# Longitudinal linked read sequencing reveals ecological and evolutionary responses of a human gut microbiome during antibiotic treatment

**DOI:** 10.1101/2019.12.21.886093

**Authors:** Morteza Roodgar, Benjamin H. Good, Nandita R. Garud, Stephen Martis, Mohan Avula, Wenyu Zhou, Samuel M. Lancaster, Hayan Lee, Afshin Babveyh, Sophia Nesamoney, Katherine S. Pollard, Michael P. Snyder

**Affiliations:** Department of Genetics, Stanford University, Stanford, California, 94305; Institute for Stem Cell Biology and Regenerative Medicine, Stanford University School of Medicine, Stanford, California, 94305; Department of Applied Physics, Stanford University, Stanford, California 94305; Department of Ecology and Evolutionary Biology, University of California Los Angeles; Department of Physics, University of California, Berkeley, CA 94720; Gladstone Institutes, San Francisco, CA 94158; Department of Epidemiology & Biostatistics, University of California, San Francisco, Ca 94158; Chan Zuckerberg Biohub, San Francisco, CA 94158

## Abstract

Gut microbial communities can respond to antibiotic perturbations by rapidly altering their taxonomic and functional composition. However, little is known about the strain-level processes that drive this collective response. Here we characterize the gut microbiome of a single individual at high temporal and genetic resolution through a period of health, disease, antibiotic treatment, and recovery. We used deep, linked-read metagenomic sequencing to track the longitudinal trajectories of thousands of single nucleotide variants within 36 species, which allowed us to contrast these genetic dynamics with the ecological fluctuations at the species level. We found that antibiotics can drive rapid shifts in the genetic composition of individual species, often involving incomplete genome-wide sweeps of pre-existing variants. These genetic changes were frequently observed in species without obvious changes in species abundance, emphasizing the importance of monitoring diversity below the species level. We also found that many sweeping variants quickly reverted to their baseline levels once antibiotic treatment had concluded, demonstrating that the ecological resilience of the microbiota can sometimes extend all the way down to the genetic level. Our results provide new insights into the population genetic forces that shape individual microbiomes on therapeutically relevant timescales, with potential implications for personalized health and disease.

## Introduction

The composition of the gut microbiome varies among human populations and individuals, and it is thought to play a key role in maintaining health and reducing susceptibility to different diseases (*1-4*). Understanding how this microbial ecosystem changes from week to week–through periods of health, disease and treatment–is important for personalized health management and design of microbiome-aware therapies (*5*).

Many studies have investigated intra-host dynamics at the species or pathway level (*6-16*). Among other findings, these studies have shown that oral antibiotics can dramatically influence the composition of the gut microbiome over a period of days, while the community often regains much of its initial composition in the weeks or months after antibiotics are removed (*7-9*). This suggests an intriguing hypothesis, in which the long-term composition of a healthy gut community is buffered against brief environmental perturbations.

However, the mechanisms that enable this ecological resilience are still poorly understood. Does species composition recover because external strains are able to recolonize the host? Or do resident strains persist in refugia and expand again once antibiotics are removed? In the latter case, do resident populations also acquire genetic differences during this time, either due to population bottlenecks or to new selection pressures that are revealed during treatment? These questions can be addressed by quantifying fine-scale microbiome genetic variation below the species or pathway level and tracking how it changes during periods of health, disease, and treatment.

Recent advances in strain-resolved metagenomics and isolate sequencing (*17-19*) have made it possible to detect DNA sequence variants within species, and to track how they change within and between hosts. These studies have shown that gut bacteria can acquire genetic differences over time even in healthy human hosts, and that these differences arise from a mixture of external replacement events (*18, 20, 21*) and the evolution of resident strains (*21-23*). However, since these previous studies have included relatively few timepoints per host, or relatively shallow sampling of their microbiota, the population genetic processes that drive these strain-level dynamics remain poorly characterized. Understanding how the forces of mutation, recombination, selection, and genetic drift operate within hosts is critical for efforts to forecast personalized responses to drugs or other therapies.

To bridge this gap, we used deep metagenomic sequencing to follow the genetic diversity within a single host microbiome at approximately weekly intervals over a period of five months, which included periods of infectious disease and the oral administration of broad-spectrum antibiotics. We used a linked-read sequencing technique to generate each of our metagenomic samples: large molecules of bacterial DNA were isolated in millions of emulsified droplets, digested into shorter fragments, and labelled with a corresponding DNA barcode to follow linked reads from the same droplet. Previous work has shown that the linkage information encoded in these barcoded “read clouds” can improve genome assembly (*24*) and taxonomic assignment (*25*) in human gut metagenomes. Here, we took a different approach and developed new statistical methods that leverage longitudinal linked-read sequencing to detect and interpret fine-scale genetic changes that take place within the resident populations of individual bacterial species over time. This reference-based strategy simultaneously captures the ecological and evolutionary dynamics of multiple strains in many resident species, without requiring assembly of complete genomes.

Here, we sought to use this approach to characterize the population genetic forces that shape native gut microbiota through periods of health, disease, antibiotic treatment and recovery. By analyzing the temporal dynamics of thousands of single nucleotide polymorphisms is 36 abundant species, we obtain new insights into the strain-level mechanisms that govern the ecological resiliency of this community, which have important potential implications for personalized health and disease.

## Results

### Longitudinal linked read sequencing of a human gut microbiome during disease and treatment

Generation of linked reads requires the preparation of long DNA fragments. We therefore developed an optimized protocol for extracting high-molecular weight (HMW) DNA from human stool samples (Methods). We used this approach to perform linked read sequencing (10X Genomics) on 19 stool samples collected from a single human subject over a period of 5 months (Fig. 1A; Supplemental Table S1). During this time, the individual was diagnosed with Lyme disease and received a two-week course of broad-spectrum oral antibiotics (doxycycline). We generated deep sequencing data for each sample (ranging from ∼8-160 Gbp), so that a typical read was present in a “read cloud” containing ∼4-30 other read pairs (Fig. 1D,E; Supplemental Fig. S1). Taxonomic profiling showed that the subject’s gut microbial community contained 66 bacterial species with median relative abundance >10^−5^, all but one of which exceeded 0.1% frequency in at least one timepoint (Fig. 1B; Methods). Consistent with previous studies (*25*), we found that the extensive taxonomic diversity of this community was accompanied by high levels of read cloud “impurity”, in which individual read clouds were frequently observed to contain fragments from ∼5-10 different species (Fig. 1E). The presence of such impurities makes it difficult to directly infer genetic linkage from individual read clouds in complex metagenomic samples. Here, we sought to overcome this issue by employing a two-stage approach, which leverages the hybrid nature of the linked-read sequencing protocol. We first ignored long-range linkage and used short-read, reference-based methods to track species and sub-species diversity over time (Methods) (*21, 22, 26*). We then developed a statistical model for linking genomic regions with higher-than-expected read cloud sharing given the observed levels of read cloud impurity (Methods). Using this hybrid approach, we documented the ecological and evolutionary responses of the gut microbial community before, during, and after antibiotic treatment.

**Figure 1:**
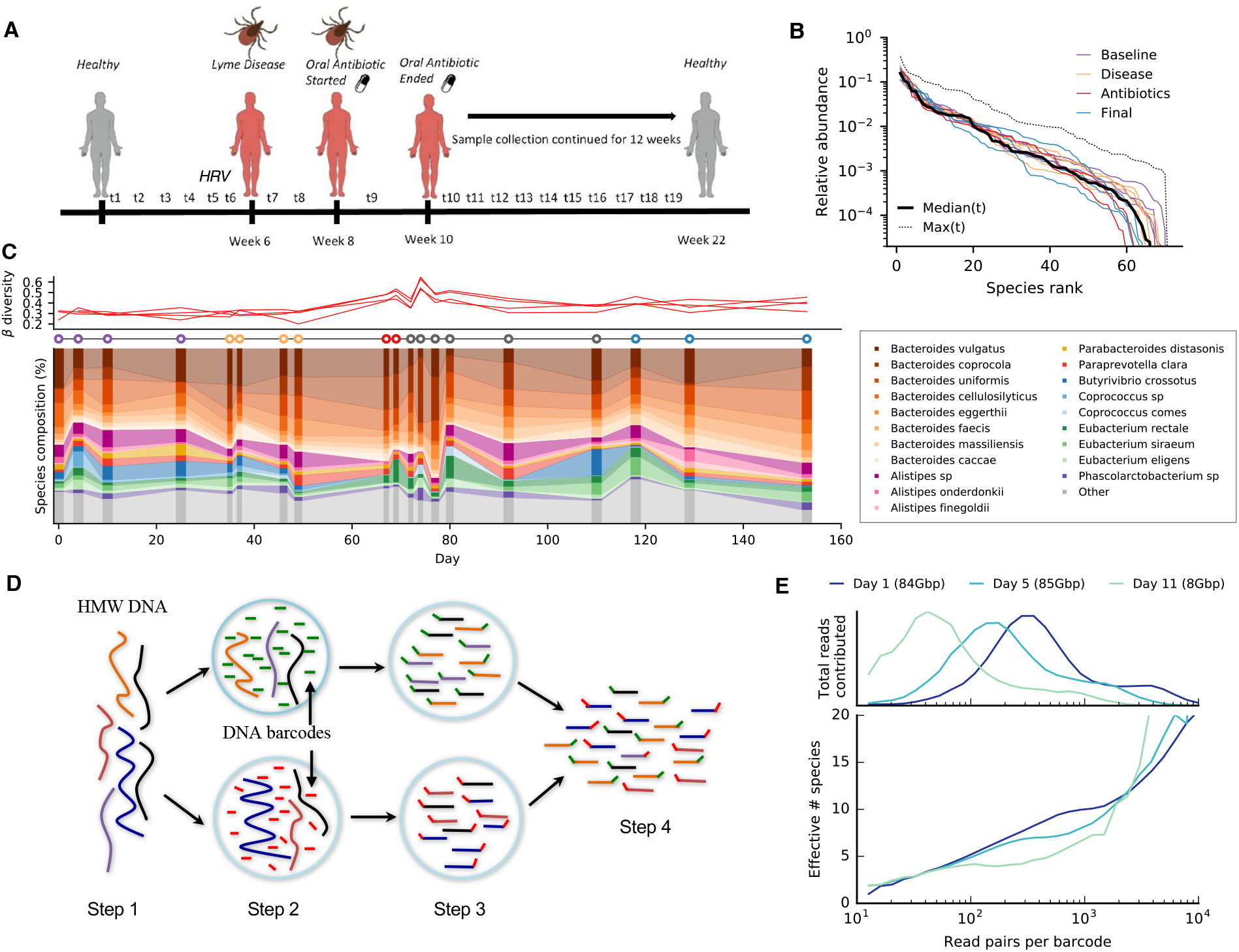
Read cloud sequencing of the gut microbiome of a single individual during disease, antibiotic treatment, and recovery. **a**, Study design. Linked read metagenomic sequencing was performed on 19 fecal samples collected from a single individual over a period of 5 months. During this time, the individual was diagnosed with Human Rhinovirus (HRV) and Lyme disease and received an oral course of doxycycline. **b**, Rank relative abundance distribution at the species level, estimated from shotgun metagenomic reads (Methods). Colored lines show distributions obtained from individual timepoints, colored according to the epochs defined in Table S1. Solid and dashed black lines denote median and maximum relative abundances across all timepoints, respectively. **c**, Species-level composition over time. Top panel illustrates Jensen-Shannon distance to each of four baseline samples as a function of time. Bottom panel shows relative abundance trajectories for a subset of the most abundant species; others are grouped together into the ‘other’ category. **d**, Schematic of linked read sequencing with the 10X Genomics platform. High molecular weight metagenomic DNA is partitioned into millions of microfluidic droplets. Amplification and ligation reactions are performed within each droplet, yielding millions of short-read libraries that are tagged with droplet-specific DNA barcodes. The resulting “read clouds” are then pooled together and sequenced on an Illumina instrument. **e**, Observed statistics of read clouds from the first three timepoints. The top panel shows total number of read pairs contributed by read clouds as a function of the number of read pairs they contain. The bottom panel shows a measure of the effective number of species that are detected in each read cloud as a function of the number of read pairs it contains (Methods). Many read clouds contain fragments from several different DNA molecules.

We first examined the ecological responses at the species level. Consistent with previous work (*20, 26-30*), we observed a substantial perturbation in species-level composition during and immediately after antibiotic treatment, followed by a return to near baseline values by the end of the study (Fig. 1C). However, we found that only a few species underwent large declines in abundance during the treatment period, and even fewer showed signs of going extinct during this interval. We used microbial DNA quantification (microbial DNA per mass of sample) (*31*) to convert our relative abundance measurements into absolute microbial densities for a subset of the study timepoints (Supplemental Fig. S2). Of the 50 species with baseline abundance >0.1%, we found that only 12 declined by more than 10-fold at the end of the two-week treatment window, and all but 4 of these recovered to near baseline values by the end of the study (Supplemental Fig. S3). *Alistipes finegoldii* and *Butyrivibrio crossotus* provide two prototypical examples of this effect: both experienced a dramatic decline in abundance during antibiotic treatment, but eventually recovered to their initial levels over the next ∼3-5 weeks (Fig. 2A). The small number of such examples, combined with the relatively steady fecal DNA content, suggests that a large fraction of the initial community was able to maintain high absolute abundance during antibiotic treatment, e.g. due to reduced sensitivity to doxycycline. Consistent with this hypothesis, we observed a high baseline proportion of doxycycline-related resistance genes among our metagenomic reads (∼200 reads per million mapped), which increased ∼2-fold during treatment (Supplemental Fig. S4). This finding is also consistent with previous observations of tetracycline resistance in isolates of several *Bacteroides* species (*32*).

**Figure 2:**
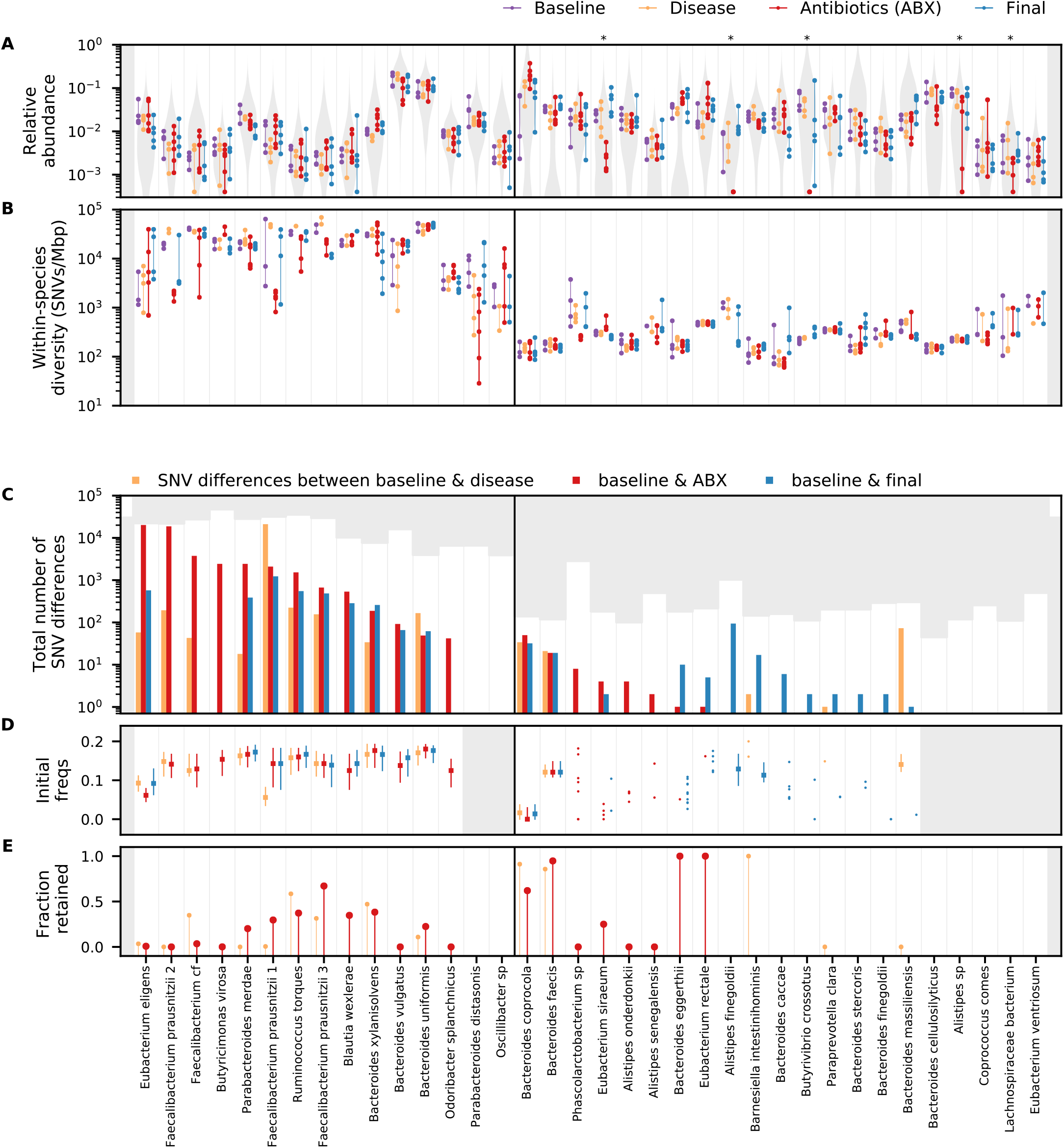
Varied ecological and genetic responses across 36 abundant species in the same host. **a**, Relative abundances of species through time, partitioned according to the epochs defined in Table S1. Each timepoint is indicated by a point, and the timepoints from the same epoch are connected by a vertical line to aid in visualization. For comparison, the grey distribution shows the corresponding values across a larger human cohort (Methods). Species whose relative abundance drops by more than 10-fold between baseline and antibiotic timepoints are indicated with a single star. Only a minority of the most abundant species experience such reductions in relative abundance during treatment. **b**, Within-species nucleotide diversity for each timepoint, as measured by the fraction of core genome sites with intermediate allele frequencies (0.2<f<0.8, Methods). Points are plotted according to the same scheme as in (a). **c**, The total number of single nucleotide (SNV) differences between a baseline timepoint and each of the later epochs (Methods). The height of the white area indicates the total number of polymorphic SNVs that were tested for temporal variation. Different species display a range of different behaviors, which can be partitioned into putative cases of competition between distantly related strains (left of vertical divider line) and evolution within a dominant resident strain (right). **d**, Initial frequencies of alleles identified in (c). For species with more than 10 SNV differences, the data are summarized by the median initial frequency (square symbol) and the interquartile range (line). Many alleles have nonzero frequency before the sweep occurs. **e**, Fraction of SNV differences in (c) that are retained at the final timepoint (f>0.7). In many species, only a minority of SNV differences gained during disease or treatment are retained.

### Deep longitudinal sequencing reveals shifts in the genetic composition of 36 species in the same host

The general pattern of persistence and recovery at the species level is shared by many other classes of antibiotics (*28*). However, the strain-level dynamics that give rise to this long-term stability remain poorly understood. Do the species that persist through disease and treatment remain stable genetically? Or does this general pattern of robustness mask a larger flux of genetic changes occurring within individual species? Our deep sequencing approach allows us to address these questions by tracking genetic variation within species over time.

We tracked the genetic composition of each species by aligning our short sequencing reads to a panel of reference genomes and estimating the population frequency of single nucleotide variants (SNVs) that we detected at each timepoint (Methods). Our high sequencing coverage enabled at least ∼10-fold coverage per timepoint for species with relative abundance >0.3%, and coverages as high as ∼500-fold in some of the most abundant species (Supplemental Fig. S5). This allowed us to simultaneously monitor SNV trajectories within 36 different species that passed our minimum coverage thresholds (Methods), and to explore how these “evolutionary” dynamics either mirrored or contrasted with the dynamics of their species abundance trajectories above (Figs. 2,3; Supplemental Figs. S6-S8; Supplemental Data 1-3).

**Figure 3:**
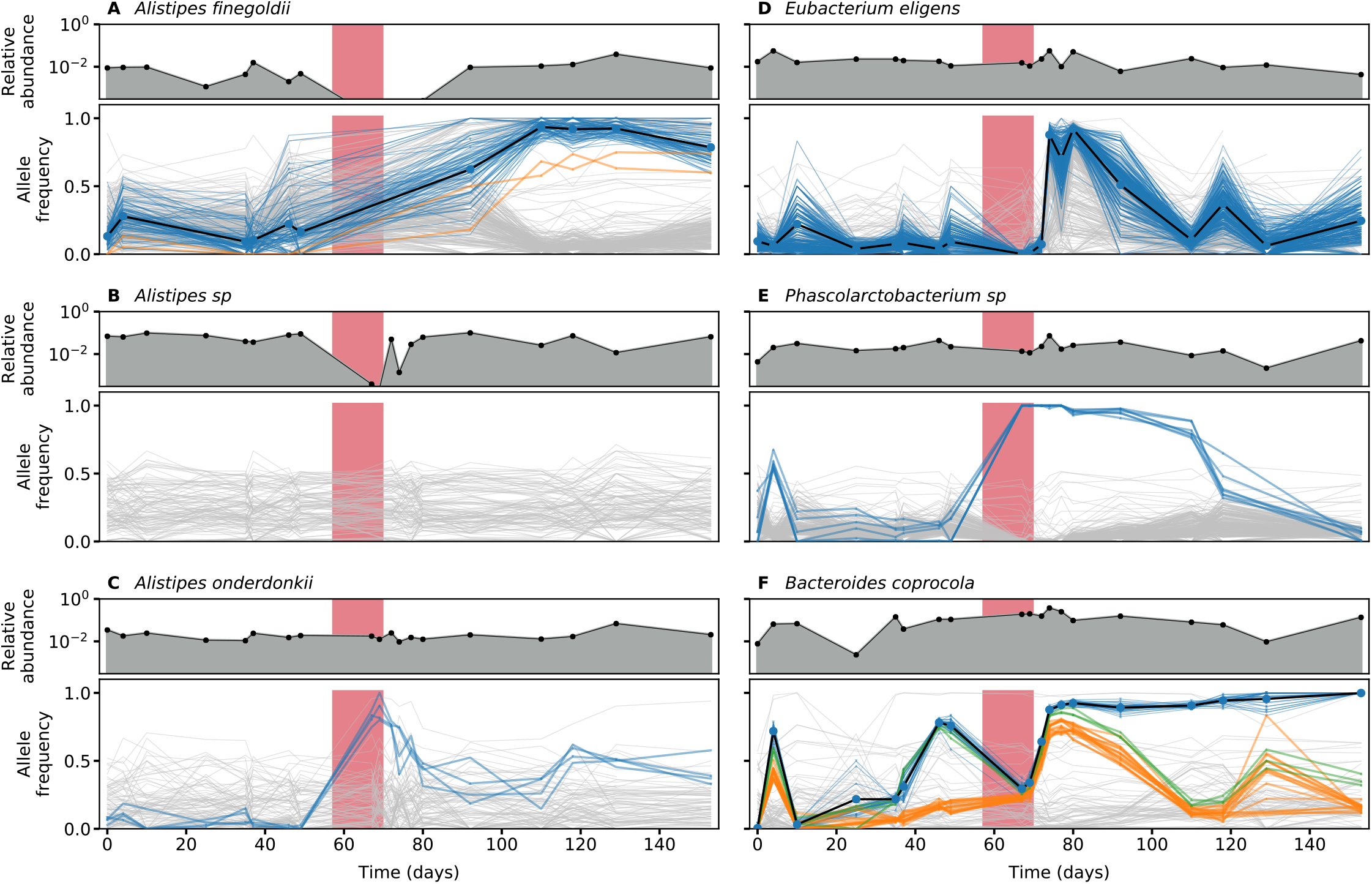
Ecological and genetic dynamics in six example species. A subset of the species in Fig. 2 were chosen to illustrate a range of characteristic behaviors (**a-f**). For each of the six species, the top panel shows the relative abundance of that species over time, while the bottom panel shows the frequencies of single nucleotide variants (SNVs) within that species. Colored lines indicate SNVs that underwent a significant shift in frequency over time (Methods), while a subset of non-significant SNVs are shown in light grey for comparison. The colors of temporally varying SNVs are assigned by a hierarchical clustering scheme, which is also used to determine their polarization (Methods).

This strain-level analysis revealed striking differences in the genetic composition of different species. Consistent with previous work (*18, 20, 21, 27*), we found that the initial levels of genetic diversity varied widely between species. Some common species, such as *Bacteroides vulgatus* and *Bacteroides uniformis*, harbored more than ∼10,000 SNVs at intermediate frequencies whereas other species, e.g. *Bacteroides coprocola* or *Alistipes sp*, had fewer than ∼100 detectable SNVs (Fig. 2B). Of particular interest are those SNVs that underwent large changes in frequency between the initial and later timepoints (e.g. from <20% to >80%, with FDR<0.1, see Methods); these “SNV differences” indicate a nearly complete “sweep” within the species of interest. We observed a similarly wide range in the number of SNV differences during and immediately after antibiotic treatment, from more than ∼10,000 in some species (e.g. *Eubacterium eligens*) to ∼10 (or even 0) in others (Fig. 2C). Of the 36 populations in Fig. 2C, more than half accumulated at least one SNV difference during this period, and more than 80% accumulated SNV differences in at least one portion of the study.

Similar within-host changes have recently been observed in metagenomic analyses from healthy hosts (*21-23*), though at a significantly lower rate (Fisher’s exact test, P<0.001). A major challenge in these earlier studies has been to demonstrate that the temporally variable SNVs are truly linked to their inferred genomic background, as opposed to read mapping artifacts (e.g., from another temporally fluctuating species that happens to share some part of the genome). Linked read sequencing provides an opportunity to address this question. For each SNV difference reported in Fig. 2C, we examined the patterns of barcode sharing with genes in the “core” genomes of our reference genome panel, which provides a proxy for the true genomic background (Methods). This analysis yielded positive confirmation for ∼80% of the SNV differences in Fig. 2C (where both alleles share read clouds with a core gene in the target species), and negative confirmation for <1% (Fig. S7). We conclude that the majority of these SNVs represent true genetic changes within their respective species populations.

The variable genetic responses in different species are not easily explained by their phylogenetic relatedness or their relative abundance trajectories. As an example, Fig. 3 shows the full species abundance and SNV frequency trajectories for six example species, which are chosen to illustrate the range of observed behaviors. This set includes three different members of the *Alistipes* genus that coexist within this particular host. The first two species, *A. sp* and *A. finegoldii*, experienced dramatic reductions in relative abundance during treatment, but we observed genetic differences in only one of the populations (*A. finegoldii*) when they recovered to their initial levels. In *A. onderdonkii*, by contrast, the relative abundance remained high at the end of the treatment phase, but we observed rapid changes in the frequencies of several SNVs within this species during the same time period (P<0.001, Methods). These examples show that species abundances alone are not sufficient to predict genetic response within species: relatively constant species abundance trajectories can mask interesting genetic shifts within a species, and vice versa.

### Quantifying genetic linkage between SNVs using barcoded read clouds

We next sought to quantify the population genetic processes that could give rise to the SNV changes observed in Figs. 2 and 3. A key question is the extent of genetic linkage within species: is recombination sufficiently frequent that genetic drift and natural selection act independently on different SNVs? Or are SNV trajectories tightly correlated because they are linked together on a small number of clonal backgrounds? This question is particularly relevant for species with high levels of SNV diversity like *B. vulgatus* (Supplemental Fig. S10), where it can mean the difference between ∼10,000 evolutionary trajectories (if SNVs are independent) or possibly only one (if SNVs derive from a mixture of two clonal strains).

Previous analyses of gut bacteria suggest that recombination can efficiently decouple SNVs over long timescales (i.e., millions of bacterial generations) (*21, 33*), but the extent of genetic linkage within hosts remains unclear. The additional information provided by linked read sequencing now allows us to investigate this question. We developed a statistical approach for detecting linkage between pairs of SNVs (Fig. 4A), which accounts for the substantial coverage variation across different species and read clouds (Methods). Fig. 4B shows how the overall levels of read cloud sharing depend on the coordinate distance between the two SNVs on the reference genome. Consistent with the fragment length estimates from our HMW DNA extraction protocol (Supplemental Fig. S11), we observed an enrichment of shared read clouds barcodes for SNVs within ∼10kb of each other, though the overall fraction of long-range read clouds remains modest (∼10%). For the subset of SNVs pairs with significant read cloud sharing, we further quantified genetic linkage by examining the combinations of major and minor alleles that are observed in the same read clouds. In particular, we estimated the number of allelic combinations (or haplotypes) that were observed for each pair of SNVs as a function of their coordinate distance along the reference genome (Fig. 4A). According to the four-gamete test (*34*), three or fewer haplotypes are consistent with clonal evolution, but the presence of all four haplotypes indicates a possible recombination event between the two SNVs (Methods). Fig. 4D shows that the vast majority of the SNV pairs we observed across species were consistent with clonal evolution: of the ∼4 million SNV pairs we examined that were separated by more than 2kb (Fig. 4C), only ∼600 showed significant evidence for all four haplotypes (q<0.05, Methods). Most of these four-haplotype pairs were concentrated in just a few species, with high values of linkage disequilibrium between the two SNVs (Supplemental Fig. S12). This suggests that to a first approximation, the SNV dynamics within species in this time course reflect a competition between a few clonal haplotypes, rather than independent alleles. This is consistent with previous indirect evidence from the clustering of allele frequencies within hosts (*21, 29, 35*).

**Figure 4:**
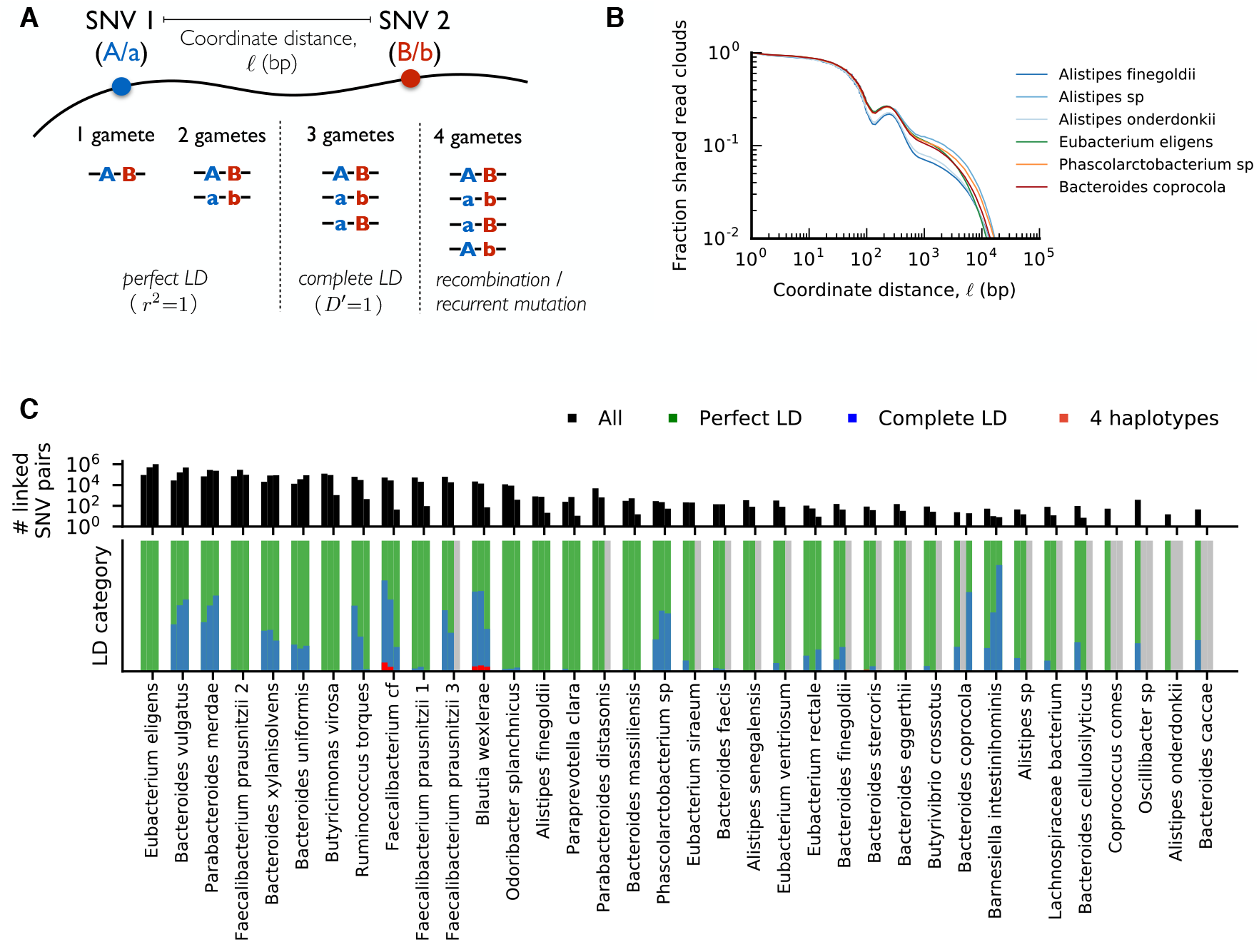
High levels of linkage disequilibrium (LD) in many resident populations. **a**, Schematic of read cloud sharing between two SNVs separated by coordinate distance *ℓ* on the same reference contig. Three or fewer haplotypes are consistent with clonal evolution, while four haplotypes indicate a possible recombination event. **b**, Observed fraction of shared read clouds as a function of *ℓ* for SNVs in the six example species in Fig. 3. **c**, Linkage disequilibrium between pairs of SNVs across a range of different species. The top panel shows the total number of linked SNV pairs (i.e., those with significantly elevated levels of read cloud sharing) for species in Fig. 2 with sufficient coverage (Methods). For each species, the three bars denote SNV pairs with *ℓ*<200bp, 200bp< *ℓ*<2kb, and *ℓ*>2kb, respectively. SNVs are included only if the minor allele has frequency f>0.1. The bottom panel shows the observed proportion of SNV pairs in the top panel that fall each of the LD categories illustrated in (a). Across species, only a small fraction of SNV pairs provide evidence for recombination.

For populations with sufficiently high SNV densities (>1 per kb), the patterns of read cloud sharing can inform efforts to cluster SNVs into smaller numbers of competing haplotypes (Supplemental Fig. S10). However, we found that many interesting temporal changes occurred in populations with much lower SNV densities (<1 per 10kb, Fig 2). In these populations, Fig. 4B suggests that SNVs will not typically share read clouds, except in rare cases where they happen to be located in nearby regions of the genome. We therefore used a heuristic approach to infer clusters of perfectly linked SNVs (a form of multi-SNV haplotype) based on similarities in their allele frequency trajectories (Methods). The inferred haplotypes are indicated by the coloring scheme in Fig. 3.

### Temporal dynamics of haplotypes reveal cryptic phenotypic differences and time-varying selection within species

We next investigated the role that natural selection plays in driving the genetic changes we observed within species. While adaptive evolution is ubiquitous in microbial evolution experiments (*36*) and many pathogens (*37, 38*), its influence on natural genetic variation in the human gut microbiota is less well understood. One common model assumes that within-host dynamics are dominated by selection at higher levels of taxonomic organization (e.g. species, families, or functional guilds), while closely related variants that survive this environmental filter are largely interchangeable. In this “conditionally neutral” scenario, short-term changes in the genetic composition of a resident population primarily occur through stochastic demographic processes like genetic drift (*39, 40*) or genetic draft (*41*). Empirical support for this model is currently mixed: one recent study in *Bacteroides fragilis* found signatures of non-neutral evolution (*23*), while a second study in *Escherichia coli* came to the opposite conclusion, potentially due to a smaller effective population size in *E. coli* (*22*). At present, it is not clear which of these scenarios will apply to the other species in Figs. 2 and 3, or how they generalize the stronger environmental perturbations associated with antibiotic treatment. These sudden environmental shifts could uncover cryptic phenotypic differences between previously coexisting strains, driving rapid shifts in the frequencies of segregating genetic variants (*42*).

High levels of genetic linkage make it difficult to distinguish these scenarios using traditional population genetic approaches, since selection and drift will typically act on extended haplotypes rather than individual alleles. For example, the sweeping SNV clusters we identified in Fig. 3 contain many synonymous variants (Fig. 5), which are most likely hitchhiking alongside causative “driver” mutations that are located in other linked regions of the genome. These driver mutations may not even be visible in Fig. 3 if they happen to arise from structural variants, mobile elements, or other accessory genes that are difficult to detect in our metagenomic pipeline (*23, 24, 43-45*).

**Figure 5:**
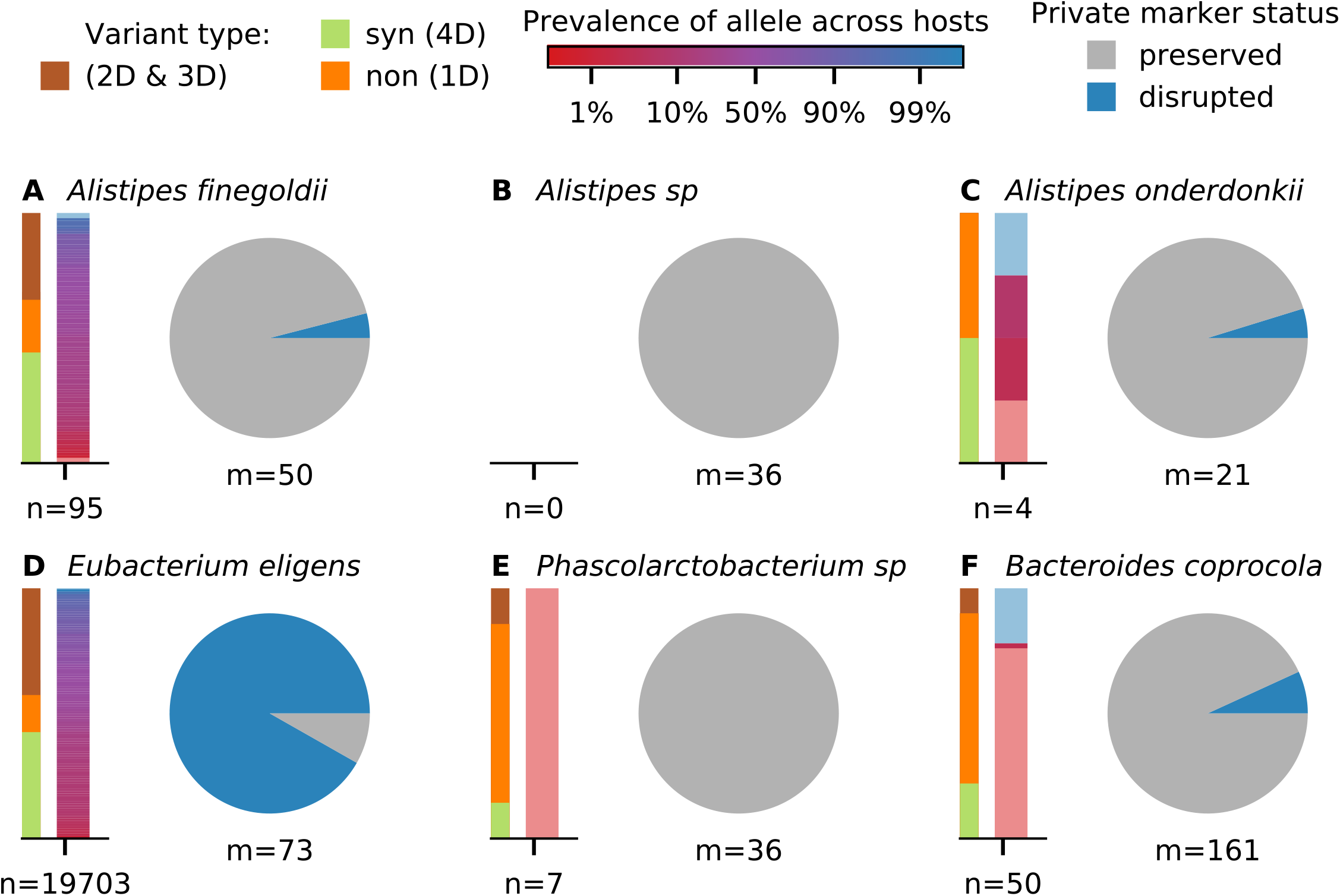
Signatures of strain replacement and evolutionary modification. **a-f**, Statistical properties of temporally varying SNVs from the six example species in Fig. 3. For each species, the bars on the left show the relative proportion of SNVs with different protein coding effects and allele prevalence across other hosts in a larger cohort (Methods). Protein coding effects are estimated from the codon degeneracy at each site (4D=fourfold degenerate/synonymous, 1D=one-fold degenerate/nonsynonymous). Allele prevalences for SNVs not observed in other hosts are indicated by light red or blue shading. Pie charts indicate the relative proportion of private marker SNVs for each species that are preserved or disrupted throughout the sampling interval (Methods). Large fractions of disrupted marker SNVs indicate a strain replacement event.

To overcome these limitations, we sought to develop a methodology for distinguishing natural selection and demographic stochasticity using the residual information encoded in the shapes of the SNV trajectories in Fig. 3. We first developed a statistical test to determine whether the SNV clusters in Fig. 3 were consistent with the simplest neutral null model, with a constant but unknown strength of genetic drift (Methods). This test leverages the fact that the frequency changes produced by a constant rate of genetic drift must be statistically similar over time, so that a large fluctuation in one time-window is unlikely to be followed by a small change in another. Our observed trajectories often violate this prediction, and allowing us to reject the constant genetic drift model for 4 of the 5 species in Fig. 3 (Supplemental Table S3).

A second possibility is that genetic drift is significantly elevated during antibiotic treatment, e.g. due to a transient population bottleneck. This could be a particularly plausible hypothesis for the *Alistipes finegoldii* population in Fig. 3A, where the genetic shift coincided with a dramatic reduction in the relative abundance of that species. While it is difficult to rule out similar bottlenecks at unobserved timepoints for the other species in Fig. 3, we still observe significant departures from the constant genetic drift model for these species even when the antibiotic timepoints are excluded (Supplemental Table S3). Closer inspection of the frequency trajectories reveals the likely source of this signal: many of the SNV clusters continue to change in frequency, but in the opposite direction, even after antibiotic treatment has concluded. This behavior, which is recapitulated across the larger set of species in Fig. 2E, cannot be explained by a simple bottleneck during treatment. Instead, these data suggest the initial increases and later reversals are most likely caused by time-varying selection pressures that act on different haplotypes within these populations, reflecting the complex environmental conditions experienced by these species *in vivo*.

The high temporal resolution of the SNV clusters allows us to estimate the magnitudes of these temporally varying selection pressures in different time windows. For example, the frequency reversals that occurred after treatment in Fig. 3 took place over ∼30-40 days, implying a fitness difference about S∼10% per day (Methods). The increases in frequency during and immediately after antibiotics were even more rapid. In *Eubacterium eligens*, the minority haplotype increased from 7% to 90% in just two days, implying a corresponding fitness difference of at least S∼250% per day. The presence of these large fitness differences is potentially not surprising, given the strong environmental perturbations that are presumably imposed by antibiotic treatment. Interestingly, however, we found that the large frequency shift in *E. eligens* did not take place during the two-week course of doxycycline, but rather 3-5 days after treatment had concluded. [For comparison, we note that serum concentrations of doxycycline usually reach their peak value after ∼4hr, with a half-life of 12-24hrs (*46*)]. We also found that the *E. eligens* population exhibited a ∼20% increase in its replication origin peak-to-trough ratio (PTR), a proxy for bacterial growth rate, during this same time interval (Supplemental Fig. S13). This indicates that the large fitness differences within this species were likely driven by a higher growth rate of the sweeping haplotype, rather than an increased death rate of the declining strain.

### Statistics of sweeping SNVs reveal strain replacement, evolutionary modification, and selection on standing variation

After demonstrating that genetic changes occur within species, and that these changes are likely driven by selection on linked haplotypes and not necessarily associated with changes in species abundance, we next sought to investigate the origin of these within-host sweeps. A key question is whether the temporally variable SNVs arose within the host or its local environment (*evolutionary modification*) or whether they reflect the invasion of pre-existing strains that diverged for many generations before colonizing the host (*strain replacement*). Following previous work (*21*), we distinguished between these two scenarios by examining three additional features of the SNV trajectories in Fig. 3: (i) the protein-coding impact of these mutations, (ii) their prevalence across other hosts in a large reference panel, and (iii) the retention of private marker SNVs (i.e., high-frequency alleles that are unique to the present host; Supplemental Fig. S14) (Methods). Figure 5 illustrates these quantities for the six example species in Fig. 3, which were chosen to cover the range of different behaviors that we observed.

The *Eubacterium eligens* population provides a striking example of strain replacement. The sweeping haplotype in this species contained more than 10,000 SNVs that were widely distributed across the genome (Supplemental Data S1), consistent with the typical genetic differences between *E. eligens* strains in different hosts (*18, 20, 21*). Few private marker SNVs were retained from the initial timepoint (Fig. 5D), which is again consistent with replacement by a distantly related strain (Supplemental Fig. S14). Similar examples of strain replacement have been observed in previous studies (*6, 10, 18, 21, 44*), but our densely sampled time course provides new information about the dynamics of this process. The SNV frequency trajectories in Fig. 3D show that the distantly related strain was already present at substantial frequencies (∼5%) long before its dramatic fitness difference was revealed. Fig. 2D shows that this is also the case for the four other putative replacement events in Fig. 2. This indicates that the replacement events we observed here were caused by the sudden increase of previously colonizing strains, rather than the contemporary invasion of new strains from outside the host.

At the opposite extreme, the *Phascolarctobacterium* population in Fig. 3E provides a prototypical illustration of an evolutionary modification event. In this case, a cluster of just 6 SNVs (including 5 amino acid changes, all in non-contiguous genes in the reference genome that are unlinked in our read clouds; Supplemental Data S1 & S4) nearly swept to fixation during antibiotic treatment (f>99.8%, S>30% per day), only to decline in frequency later in the study. Unlike the replacement event above, this sweep shared all 42 of the private marker SNVs from the dominant strain at the initial timepoint (Fig. 5E), suggesting that they recently descended from a common ancestor. Interestingly, however, we again observe evidence that the minority haplotype was already segregating at substantial frequencies (∼1-10%) before treatment, a finding which is recapitulated for several other non-replacement examples in Fig. 2D. This suggests that frequency-dependent selection may have initially driven these mutant lineages to intermediate frequencies – and maintained them there – before antibiotics or other environmental changes (or subsequent mutations) caused them to sweep through the rest of the population.

In addition to these extreme cases, we also observed a third category of events that seem to bridge the divide between strain replacement and evolutionary modification. For example, in the *Alistipes finegoldii* population in Figs. 3A and 5A, a cluster of ∼80 SNVs swept to high frequency when the species recovered from antibiotic treatment, potentially consistent with a population bottleneck. While the high rates of private marker SNVs sharing (52/55) suggest that the sweeping haplotype is a modification of the dominant strain from the initial timepoint, the large fraction of synonymous mutations (dN/dS=0.16), many of which are shared across other hosts, is more consistent with a strain replacement event. Moreover, in contrast to the two examples above, we found that the *A. finegoldii* SNVs fell into a smaller number of contiguous genes in the reference genome and were often linked together in the same read clouds (Supplemental Data S1 & S4). These same SNVs are also frequently co-inherited in “haplotype blocks” among the other hosts in our larger reference panel (Supplemental Fig. S15). Taken together, these independent lines of evidence suggest that the *A. finegoldii* SNVs in Fig. 3A were most likely transferred onto their current genetic background through recombination. Similar to the *E. eligens* and *Phascolarctobacterium* examples above, the sweeping haplotype in *A. finegoldii* was already segregating as a minor variant (f∼20%) before antibiotic treatment, suggesting that the original recombination event (and its initial rise to observable frequencies) predated the current sampling period. In fact, the same cluster of SNVs can be observed in a previously sequenced sample from the same subject that was collected roughly two years earlier (Supplemental Fig. S6; (*27*)), suggesting that the recombinant haplotype is at least several years old.

The *Bacteroides coprocola* population in Figs. 3F and 5F provides another interesting example. In this case, a cluster of 37 SNVs (including reversions of 11 of the 167 private marker SNVs) was already in the process of sweeping through the population before antibiotic treatment began. In this case, however, the sweeping mutations were scattered across many non-contiguous genes in the *B. coprocola* reference genome and are seldom observed in other hosts (Supplemental Data S1), so recombination no longer provides a parsimonious explanation. The fraction of synonymous mutations (dN/dS=0.7) also lies somewhere between the typical between-host values (dN/dS∼0.1) and within-host hitchhiking (dN/dS>=1). This suggests that the lineages may have coexisted with each other for a much longer period of time.

## Discussion

The response of gut microbial communities to antibiotics plays a crucial role in their susceptibility to pathogens (*7, 47, 48*), the spread of antibiotic resistance genes (*49, 50*), and their long-term stability (*8, 51, 52*). Numerous studies have documented the resilience of these communities at the taxonomic or pathway level (*7-16*). Yet the strain-level dynamics that give rise to this ecological robustness remain poorly characterized.

In this study, we sought to characterize these within-species dynamics by following the gut microbiome of a single individual through a period of health, disease, and the oral administration of doxycycline. We used linked read metagenomic sequencing to track the dynamics of single nucleotide variants within 36 different species, and to compare these dynamics with the broader ecological shifts at the species level. In contrast to previous applications of linked read sequencing, which have mostly focused on genome assembly (*24*) or taxonomic assignment (*25*), we developed population genetic approaches to quantify the fine-scale genetic changes that occur in these populations over time.

Consistent with our expectations, we found that antibiotic perturbations can lead to widespread shifts in the genetic composition of individual species, and at a higher overall rate than observed in healthy hosts (*21-23*). However, these within species changes were rarely consistent with the traditional picture of extinction and subsequent recolonization after treatment. Instead, we found that the genetic responses varied widely across species, with some resident populations acquiring thousands of consensus sequence differences, and others acquiring only a handful. These genetic changes were frequently observed in species without large changes in relative abundance in the sampled timepoints, and conversely, large abundance fluctuations were not always accompanied by widespread genetic changes. Furthermore, we found that some of the most dramatic fluctuations at both the species and SNV levels occurred in the weeks after treatment had concluded. Together, these findings suggest that the response to doxycycline is not driven by discrete recolonization events, but rather, by the subtler processes of strain-level competition and evolution within the host.

At this population genetic level, our observations revealed qualitative departures from the simplest models of neutral evolution or the spread of antibiotic resistance phenotypes via classic selective sweeps. Instead, the observed genetic responses were much more dynamic: we often observed partial genome-wide sweeps containing multiple linked genetic variants, many of which were previously segregating at observable frequencies before the onset of treatment. Although their frequencies increased dramatically on daily or weekly timescales, few of these variants ever fixed in their respective populations. Instead, we observed frequent reversion of sweeps at the single base pair level, consistent with temporally varying selection pressures and strong pleiotropic tradeoffs. However, these reversions rarely ended in extinction, and more commonly stabilized close to their initial pre-treatment frequency. These dynamics suggest that the sweeping haplotypes may have been stably maintained in their respective populations over time, e.g. due to metabolic or spatial niches. This provides a potential explanation for the “oligo-colonization” structure observed in a variety of within-host microbial populations (*18, 21, 53*). Interestingly, our data show that similar dynamics can occur for mixtures of distantly related strains, as well as for haplotypes that likely evolved within the host. This suggests that ongoing ecological diversification could play an important role in shaping the genetic structure of resident populations, echoing a previous finding in *Bacteroides fragilis (23)* .

There are several important limitations to our study. First, since we have focused primarily on single nucleotide variants in well-behaved regions of reference genomes, we are likely missing many of the true targets of selection, particularly in the case of antibiotic resistance where mobile elements (*45, 54*) and other structural rearrangements (*43*) are known to play an important role. This makes it difficult to know what fraction of genetic changes are a direct response to antibiotics, as opposed to indirect responses produced by fluctuations in the abundances of other species. It is even possible that nearly all of the mutations that we observed are simply passenger mutations that are hitchhiking alongside the true causative variants. The situation could potentially be improved by combining our read mapping approach with *de novo* genome assembly, similar to previous studies (*44, 55, 56*), particularly if linked reads are used during the assembly step (*24*). However, given the high levels of genetic linkage we inferred, it would be difficult to pinpoint individual selection pressures even with an exhaustive list of mutations, since one can only observe the net effects of selection across entire haplotypes. Our current reference-based approach is effectively using this limitation to our advantage, by relying on the dynamics of linked passengers to provide information about the net selection pressures on their corresponding haplotypes.

Our analysis also revealed potential limitations of linked read sequencing for studying strain-level dynamics in the gut microbiota. While the longer effective read lengths allowed us to resolve genetic linkage in some species or genomic regions that happened to contain high-levels of intra-host diversity, we found that many interesting temporal dynamics occurred among haplotypes with only a few SNV differences between them (≪1 per 10kb), which would be difficult to capture with typical ∼10kb fragments. This length scale also poses challenges for linking shared or accessory regions with their corresponding species “backbone”, since transferred DNA segments and other structural variants are expected to have a similar length scales. This suggests that alternative approaches like chromatin conformation capture (Hi-C) (*44, 57*) or linked-read sequencing of single cells (*58*) might offer advantages over traditional long-read sequencing for resolving the dynamics of longer range forms of genetic linkage.

In addition to these methodological constraints, a second key limitation is our focus on a single host microbiome. While the concentrated resources allowed us to observe a variety of different responses across individual species in the same community, further work will be required to establish the prevalence of these different patterns across larger cohorts, and among different classes of antibiotics. Our high-resolution time course provides a valuable set of templates that can inform future classification efforts in larger, but lower resolution studies.

In summary, by tracking a host microbiome through periods of disease, antibiotic treatment, and recovery, we uncovered new evidence that the ecological resilience of microbial communities might extend all the way down to the genetic level. Understanding how this resilience arises from the complex interplay between host genetic, epigenetic, and lifestyle factors, as well its implications for broader evolution of the microbiome, remains an exciting avenue for future work.

## Methods

### Sample Collection and Sequencing

Stool samples were collected from a 62-year-old male over a 5-month period and stored on dry ice immediately after collection (Supplemental Table S1; Supplemental Materials, Section 1). Approximately ∼200mg of stool were aliquoted from each sample for HMW DNA extraction. HMW DNA was extracted using QIAamp DNA Stool Mini Kit (Qiagen) and quantified using Qubit assay (ThermoFisher Inc., Supplemental Data S5 & S6; Supplemental Materials, Section 2). Barcoded sequencing libraries were prepared using the 10X Genomics Chromium™ platform (10X Genomics) and were sequenced using an Illumina HiSeq4000 machine (Illumina) with a read length of 2×151bp and a depth of 30-500 million reads per sample. After sequencing, 10X barcode strings were extracted from each read, and reads were subsequently trimmed to remove low quality sequence in the first 20-25bp (Supplemental Materials, Section 3). Reads were assigned to read clouds using a lightweight error correction algorithm, which merges barcode strings that are a single edit away from the inferred read cloud (Supplemental Materials, Section 4).

### Estimating species abundances

Species abundances were estimated using the MIDAS software package (*26*), which maps short sequencing reads against a database of universal single-copy marker genes from ∼6000 bacterial species (Supplemental Materials, Section 5.1). The relative abundance of each species is estimated from the relative coverage of its associated marker genes. The reference distributions in Fig 2 were obtained by applying this same approach to a panel of ∼900 healthy human fecal metagenomes collated in our previous study (*21*). Absolute abundances were estimated for a subset of the study timepoints using microbial DNA quantification (total DNA per mass of sample) (*31*). The absolute abundance of each species was estimated by scaling its relative abundance by the total concentration of extracted DNA (Supplemental Fig. S2), which serves as a proxy for the total microbiome density in that sample (*59*).

### Empirical estimates of read cloud impurity

The read cloud impurity estimates in Fig. 1E were obtained using a custom version of the MIDAS software package (*26*), which was extended to track the read cloud labels associated with each short sequencing read (Supplemental Materials, Section 5). Short sequencing reads were mapped against a curated database of gene families (or “pan-genome”) constructed from the subset of species that were detected above a minimum abundance threshold in at least one timepoint (Supplemental Materials, Sections 5.1 and 5.2). For each read cloud μ, we calculated the total number of associated short reads (*r*_μ,s_) that mapped to each species s in the pan-genome database. The effective number of species (*S*_μ_) was approximated by the root mean squared estimator,

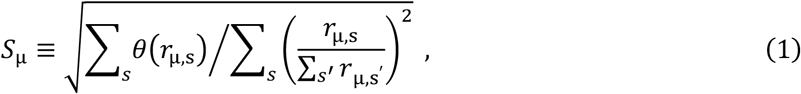

where *θ*(*z*) is the Heaviside step function (Supplemental Materials, Section 7.1). The curves in Fig. 1E were obtained by binning read clouds by their total coverage and plotting the median value of *S*_μ_ for each bin.

### Identifying SNVs within species

Single nucleotide variants (SNVs) were identified using a custom version of the pipeline developed in our previous study (*21*), which is based on the MIDAS software package (*26*). Short sequencing reads were mapped against a curated panel of reference genomes constructed from the subset of species that were detected above a minimum abundance threshold in at least one timepoint (Supplemental Materials, Sections 5.1 and 5.3). After applying a sequence of coverage filters, single nucleotide variants (SNVs) were identified from the read pileups at all remaining protein coding sites (Supplemental Materials, Sections 5.2 and 5.3). The reference allele for each SNV was chosen to coincide with the consensus allele observed within a larger human cohort that we analyzed in our previous study (*21*). This cohort was also used to estimate the prevalence of each allele within the broader human population, which we defined as the fraction of metagenomic samples in which the target allele was present in a majority of the reads (Supplemental Materials, Section 5.3.1). Following re-polarization, SNVs were retained for downstream analysis if the alternate allele was present with at least 10% frequency in at least one timepoint. Note that this definition also includes sites that are fixed within the focal host, but where the host-wide consensus is a minority allele within the larger human cohort; these sites are important for the private marker SNV analysis described below. The detected SNVs were used to estimate the overall levels of genetic diversity in Fig. 2A, defined as the fraction of sites in the “core genome” (Supplemental Materials, Section 5.2) in which the alternate allele was present at an intermediate frequency (0.2<f<0.8) (Supplemental Materials, Section 5.3.1).

Allele frequency trajectories were estimated for each SNV based on the number of unique read clouds that supported each allele. We developed an extension of the MIDAS software package (*26*) to track the read cloud labels associated with a subset of the SNVs identified above, leveraging the additional degeneracy within a read cloud to automatically correct for some sequencing and PCR errors (Supplemental Materials, Section 5.4). After applying a second round of coverage filters, we estimated the allele frequency at site *i* and timepoint *t* using the plug-in estimator 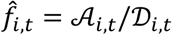, where 𝒜_*i,t*_ is the number of unique read clouds supporting the alternate allele, and 𝒟_*i,t*_ is the total number of unique read clouds that mapped to that site (Supplemental Materials, Section 5.5).

### Identifying SNV differences over time

We used the allele frequency trajectories above to identify subsets of SNVs that experienced large changes in frequency between pairs of sequenced timepoints (Supplemental Materials, Section 5.5.1). We refer to these as “SNV differences,” since they indicate a full or partial “sweep” of the allele through the resident population of interest. For each ordered pair of timepoints, we first identified all SNVs whose minor allele frequency transitioned from <20% frequency in the initial timepoint to >70% frequency in the latter timepoint. We then compared the observed number of SNVs that satisfied this criterion with the expected number under a null model of binomial sampling noise, which yields an associated P-value and false discovery rate (FDR) for the ensemble of putative SNV differences between that pair of timepoints. All SNVs in the ensemble were declared to be SNV differences if the P-values and false discovery rates exceeded a desired significance threshold (P<0.05 and FDR<0.1). Additional details about the detection algorithm are provided in Supplemental Materials, Section 5.5.1.

### Linking SNV differences with their inferred species backbone

We verified that SNV differences were linked to their inferred species backbone by examining patterns of read cloud sharing with the other reference genomes in our curated panel above (Supplemental Fig. S9). For each identified SNV difference in Fig. 2C, we created a list of the core genes in each species that shared read clouds with either of the two alleles, and we calculated the fraction of frequently shared genes that originated from the correct species (Supplemental Materials, Section 7.5). We considered there to be a positive confirmation if, for both alleles, more than two thirds of the frequently shared genes originated from the correct species. Conversely, we considered there to be a negative confirmation if either of the alleles had more than a third of their frequently shared genes originating from a different species.

### Quantifying genetic linkage between pairs of SNVs

We quantified genetic linkage between pairs of SNVs based on the distribution of shared read clouds among their four combinations of alleles (or “haplotypes”). This task is complicated by the high levels of read cloud impurity in Fig. 1E: since a typical read cloud contains several fragments, individual examples of read cloud sharing do not provide conclusive evidence for genetic linkage on their own. We therefore adopted a statistical approach, which aims to quantify elevated rates of read cloud sharing above those expected from read cloud impurities alone (Supplemental Materials, Sections 7.3 & 7.4). We first compiled a list of SNV pairs within each species with at least 12 shared read clouds across all study timepoints. We then compared the observed number of shared read clouds for each pair with a null model of read cloud impurity, in which the probability that SNVs *i* and *j* share read cloud *μ* by chance is given by

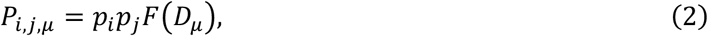

where *p*_*i*_ and *p*_*j*_ are related to the relative abundances of the SNVs at that timepoint, and *F*(*D*_*μ*_) is a function that depends on the total number of reads that were sequenced from that read cloud (Supplemental Materials, Section 7.3). We estimated these parameters empirically using the observed rates of read cloud sharing between species, under the assumption that the vast majority of this sharing was caused by read cloud impurities. We then used this model to obtain an associated P-value for each pair of SNVs above, which quantifies how their observed levels of barcode sharing differ from those expected under read cloud impurities alone (Supplemental Materials, Section 7.4.1). The top panel in Fig. 4C shows the total number of SNV pairs in each species whose Bonferroni corrected P-values were <0.1. For each of these significantly linked SNV pairs, we then repeated this calculation at the allelic level to quantify read cloud sharing among the four possible combinations of alleles for each SNV pair (Supplemental Materials, Section 7.4.2). Deviations from the read cloud impurity null model were used to classify SNVs into the three LD categories in Fig. 4. Further details on the statistical model and genetic linkage classifications are provided in Supplemental Materials, Sections 7.3 & 7.4.

### Inferring clusters of linked SNVs from correlated allele frequency trajectories

The SNV clusters in Fig. 3 were inferred using a heuristic approach that leverages correlations in the underlying allele frequency trajectories (Supplemental Materials, Section 8). In contrast to previous strain detection methods that aim to resolve the full set of haplotypes within a population (*29, 35, 56, 60*), we employed a lightweight approach that only identifies clusters of SNVs that are likely to be perfectly linked to each other, while also accounting for uncertainty in their relative polarization. For each pair of SNVs *i* and *j*, we defined a pair of dissimilarity values,

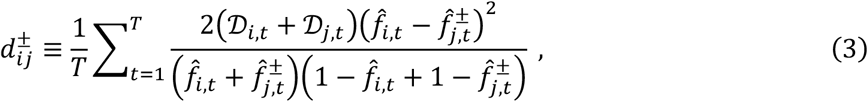

where 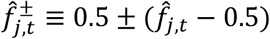 denotes the two possible polarizations of SNV *j*. We performed UPGMA clustering on the dissimilarity matrix 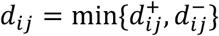, and we used a maximum distance criterion with an empirically derived threshold (*d*^*\**^ ≈ 3.5) to identify flat SNV clusters from the resulting UPGMA dendrogram. The relative values of 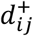 and 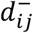 were used to determine the relative polarizations of the SNVs in each cluster, and an average frequency trajectory was estimated for the cluster by summing the ancestral and derived read counts for each of these re-polarized SNVs. Further details of the clustering algorithm, as well as comparisons to other strain detection methods, are described in Supplemental Materials, Section 8.

### Inferring natural selection and genetic drift from haplotype frequency trajectories

We sought to quantify the relative contributions of natural selection and genetic drift on the SNV differences in Fig. 3 by comparing the observed trajectories to a population genetic model for the underlying haplotype frequencies,

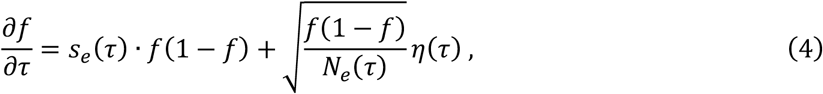

where *η*(*τ*) is a Brownian noise term(*61*), and *s*_*e*_(*τ*) and *N*_*e*_(*τ*) are effective parameters that represent the aggregate effects of natural selection and genetic drift, respectively (Supplemental Materials, Section 9). The observed frequencies 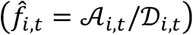 are generated from this underlying trajectory via an additional sampling step,

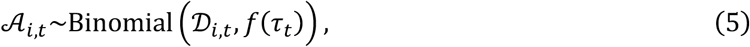

which models the finite sampling noise that occurs during sequencing. Eqs (4) and (5) constitute a standard Hidden Markov Model (HMM) for modeling genetic timeseries (*62-65*). However, this model differs from previous approaches in that we explicitly allow for time-varying selection coefficients, which are necessary to account for potential environmental heterogeneity (e.g. due to antibiotics) or linkage with other selected mutations.

In the absence of genetic drift [*N*_*e*_(*τ*) · *s*_*e*_(*τ*) · *f*(*τ*)(1 − *f*(*τ*)) ≫ 1], these time-varying selection pressures can be inferred from the plug-in estimator,

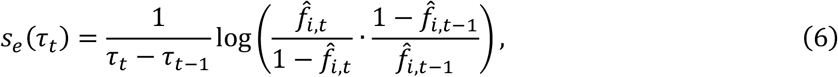

which by definition perfectly reproduces the observed frequency trajectory. We therefore asked whether this same trajectory could have plausibly emerged from genetic drift and/or sequencing noise alone.

We first developed a statistical test to determine whether the observed mutation trajectory was consistent with the simplest neutral null model, with a constant but unknown strength of genetic drift (Supplemental Materials, Section 9). Briefly, we first used dynamic programming to calculate the likelihood of the observed trajectory, Λ(*N*_*e*_), across a grid of *N*_*e*_ values to infer the maximum likelihood estimator,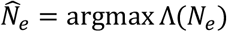. We then used the maximum likelihood 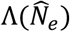 as a test statistic, and performed parametric bootstrapping using simulated trajectories to calculate an associated P-value for the observed trajectory under the null hypothesis. This test was performed for each of the inferred SNV clusters in Fig. 3, using the average frequency trajectory inferred via the clustering algorithm above (Supplemental Table S3). We also performed a generalization of this test with a variable *N*_*e*_(*τ*) in which we inferred separate *N*_*e*_ values for the segments of the trajectory that occurred before, during, and after antibiotics. This allowed us to simulate the effects of a simple bottleneck during treatment. Additional details and mathematical definitions are provided in Supplemental Materials, Section 9.

### Quantifying retention of private marker SNVs over time

Following previous work (*21, 26, 66-68*), we sought to distinguish instances of strain replacement and evolutionary modification by examining the retention of so-called “private marker SNVs” before, during, and after the sweep (Supplemental Fig. S14; Supplemental Materials, Section 5.5.2). We used a generalization of the approach we developed in our previous study (*21*) for analyzing private marker SNV sharing between pairs of sequenced timepoints. We considered a SNV to be “private” if the prevalence of its alternate or reference allele (as defined in Supplemental Materials, Section 5.3.3) was equal to zero. We then defined the set of “disrupted private marker SNVs” to be the subset of temporally variable SNVs whose reference alleles (as shown in Fig. 3) are private; the definition of a temporally variable SNV implies that these private alleles must lose their majority status at some point during the time course. Conversely, we defined the set of “preserved private marker SNVs” to be the subset of private SNVs in which the private allele remained at high frequency in a large fraction of the timepoints (specifically, >80% frequency in at least 80% of the timepoints, and no lower than 50% frequency in any one timepoint). These definitions were used to construct the pie charts in Fig. 5. Additional details and motivation for this approach are provided in Supplemental Materials, 5.5.2.

### DATA ACCESS

Raw sequencing reads have been deposited in the NCBI BioProject database under accession number PRJNA680579. All associated metadata, as well as the source code for the sequencing pipeline, downstream analyses, and figure generation, are available at GitHub (https://github.com/bgoodlab/highres_microbiome_timecourse).

## ACKNOWLEDGEMENTS

We thank Eitan Yaffe for comments on the manuscript. This work was funded in part by the US National Institutes of Health grants U54DK10255603, R01AT01023202, and 2RM1HG00773506. Sequencing was performed at the Stanford Center for Genomics and Personalized Medicine supported by US National Institutes of Health grant S10OD020141. N.R.G. and K.S.P. acknowledge support from the US National Science Foundation (DMS-1563159), the Chan Zuckerberg Biohub, and the Gladstone Institutes. B.H.G. acknowledges support from the Miller Institute for Basic Research in Science and a Stanford Terman Fellowship.

## AUTHOR CONTRIBUTIONS

M.R. and M.P.S. conceived the study; B.H.G. designed the analysis; M.R. and M.A. developed the HMW DNA extraction protocol and performed the experiments; S.L., H.L., A.B., M.R., S.N., and W.Z. performed sequencing QC and preliminary bioinformatic analyses; B.H.G., N.R.G, and S.M. developed the metagenomic pipeline and analyzed SNV data; B.H.G., S.M., and K.S.P. developed theory and statistical methods; K.S.P. and M.P.S. supervised the study; M.R. and B.H.G. wrote the paper; M.R., B.H.G., N.R.G., K.S.P., and M.P.S. edited the paper. Correspondence and requests for materials should be addressed to B.H.G., K.S.P., or M.P.S.

## SUPPLEMENTAL MATERIALS

Supplemental Methods, Supplemental Figures S1-S14, Supplemental Tables S1-S3, Supplemental Data S1-S7.

