## Supplemental Figures and Methods for "Longitudinal linked read sequencing reveals ecological and evolutionary responses of a human gut microbiome during antibiotic treatment"

### Supplemental Materials

#### Contents

Page

**Experimental Methods** **17**

**1 Study design and sample collection** **17**

**2 High molecular weight DNA extraction** **17**

**3 Linked-read library preparation and sequencing** **18**

**4 Postprocessing and read cloud assignment** **18**

**Computational Methods and Analysis** **19**

**5 Metagenomic pipeline** **19**

**8 Clustering SNVs into haplotypes** **32**

**9 Distinguishing between genetic drift and natural selection in allele frequency time series** **35**

**Supplemental Data Files** **40**

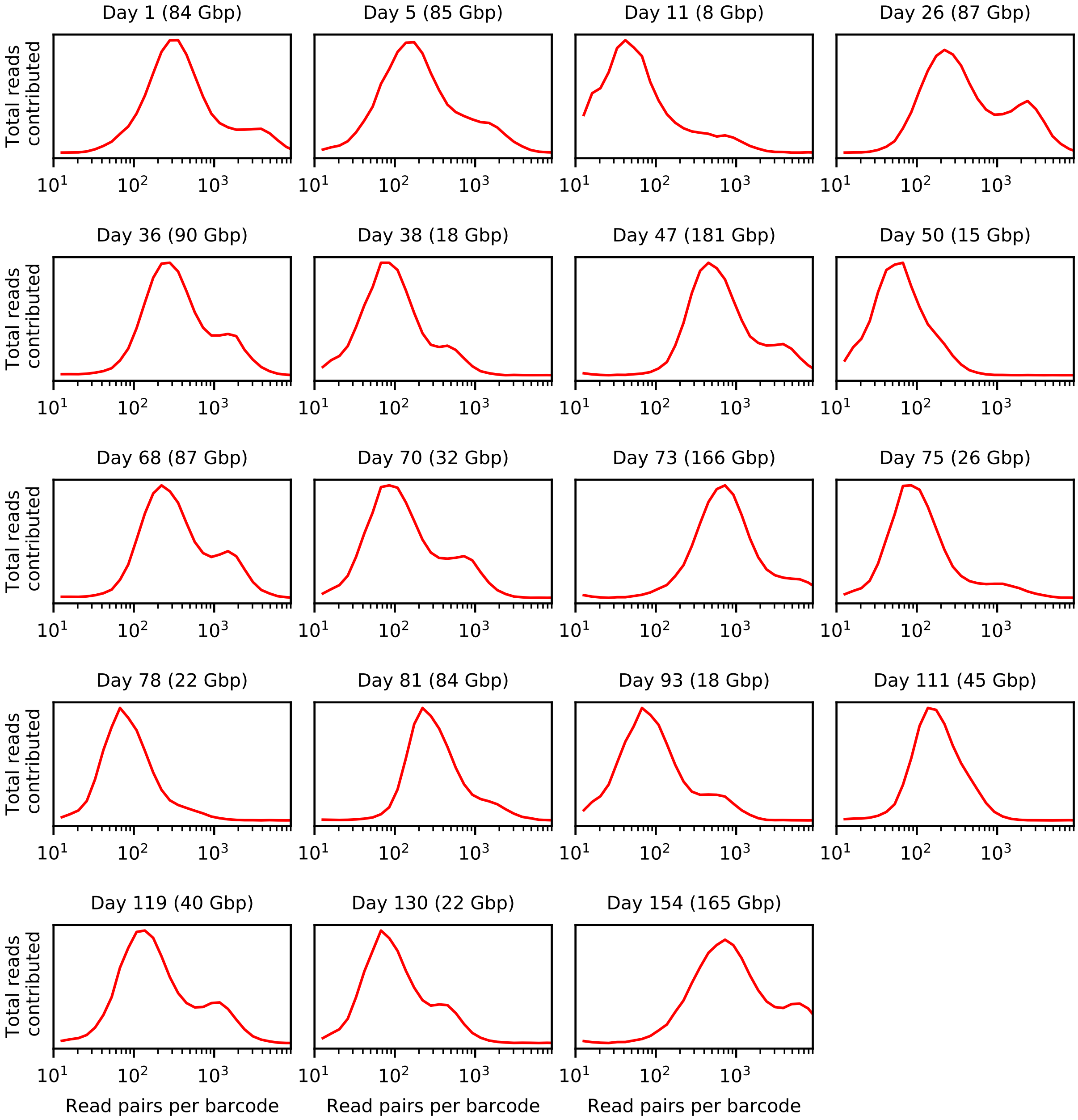

Figure S1: **Read cloud statistics for each sample.** Analogous versions of Fig. 1E for each of the 19 samples.

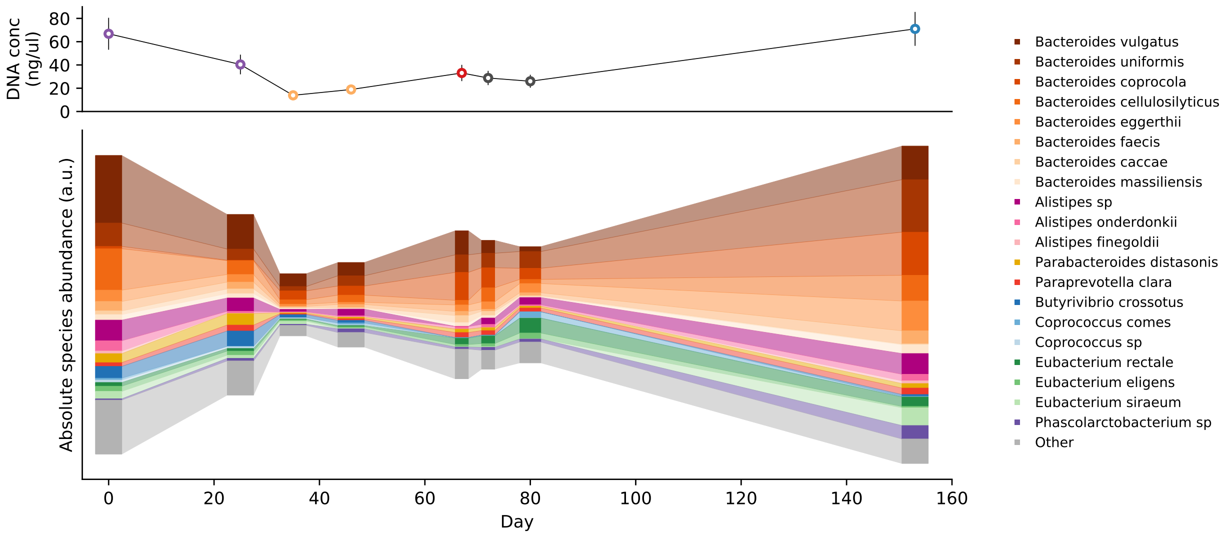

Figure S2: **Microbiota density over time.** Top: Microbiota density estimated by microbial DNA quantification (concentration of extracted DNA per ∼200mg feces) for a subset of the study timepoints, labeled using the same color scheme as in Fig. 1. Vertical lines indicate ±20% error bars, as estimated from the precision of our sample mass measurements. These data show that overall microbiota density was not significantly reduced at the end of antibiotic treatment. Bottom: Absolute species abundance dynamics obtained by scaling the relative abundance measurements in Fig. 1C by the microbiota density measurements in the top panel.

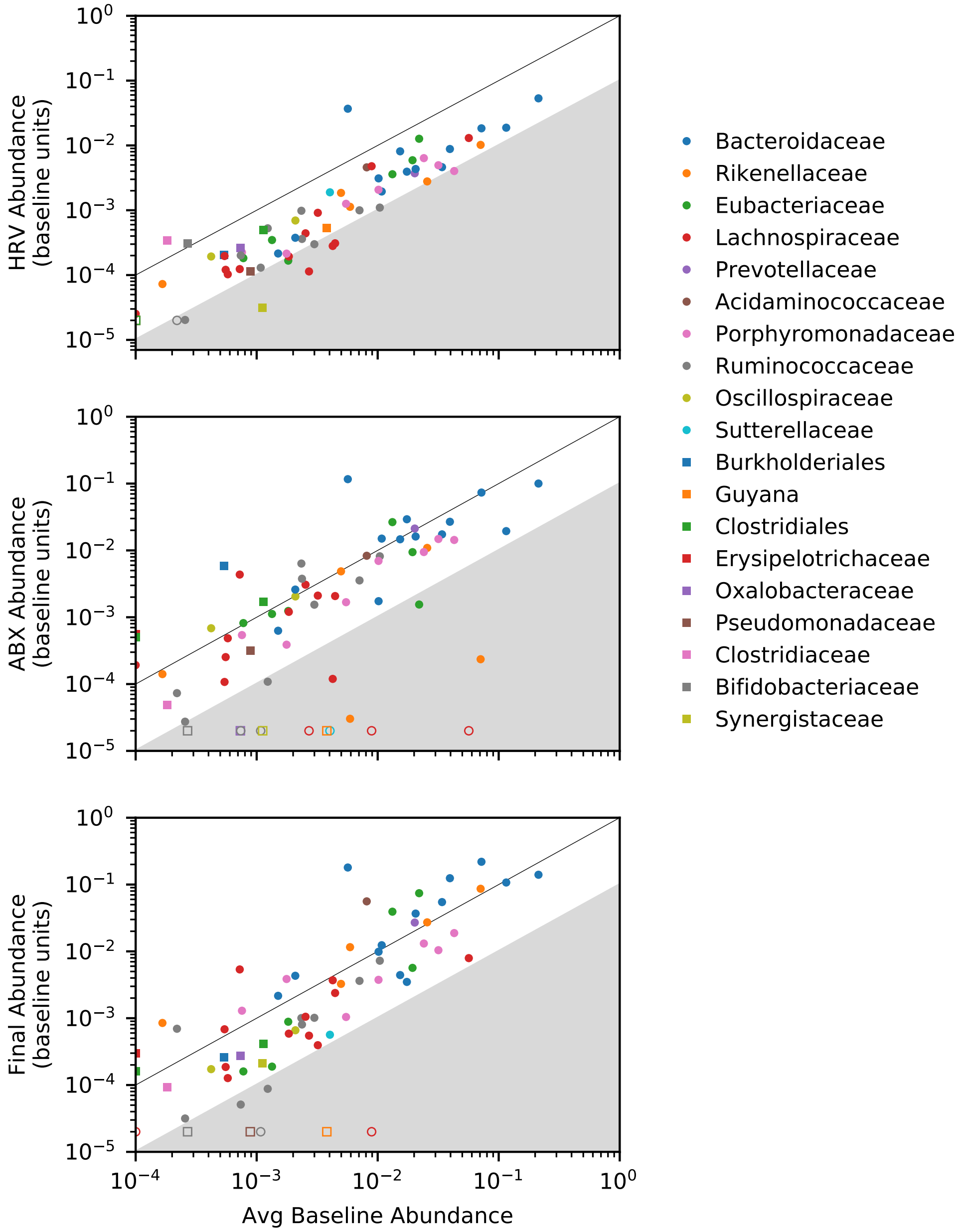

Figure S3: **Species abundance dynamics before, during, and after antibiotics.** Absolute abundances of individual species at the HRV (top), antibiotic (middle), and final timepoints (bottom) as a function of the average absolute abundance during the two baseline timepoints in Supplemental Fig. S2. Absolute abundances are estimated by multiplying relative abundance estimates by the microbial density measurements in Supplemental Fig. S2, and scaling by the average microbial density at the two baseline timepoints. Symbols denote individual species and are colored by taxnomic family. Open symbols denote species that fall below the detection threshold ($\approx 2 \times 10^{-5}$). These data show that only a minority of species experienced dramatic reductions in absolute abundance by the end of antibiotic treatment; several of these species recovered in abundance by the end of the study.

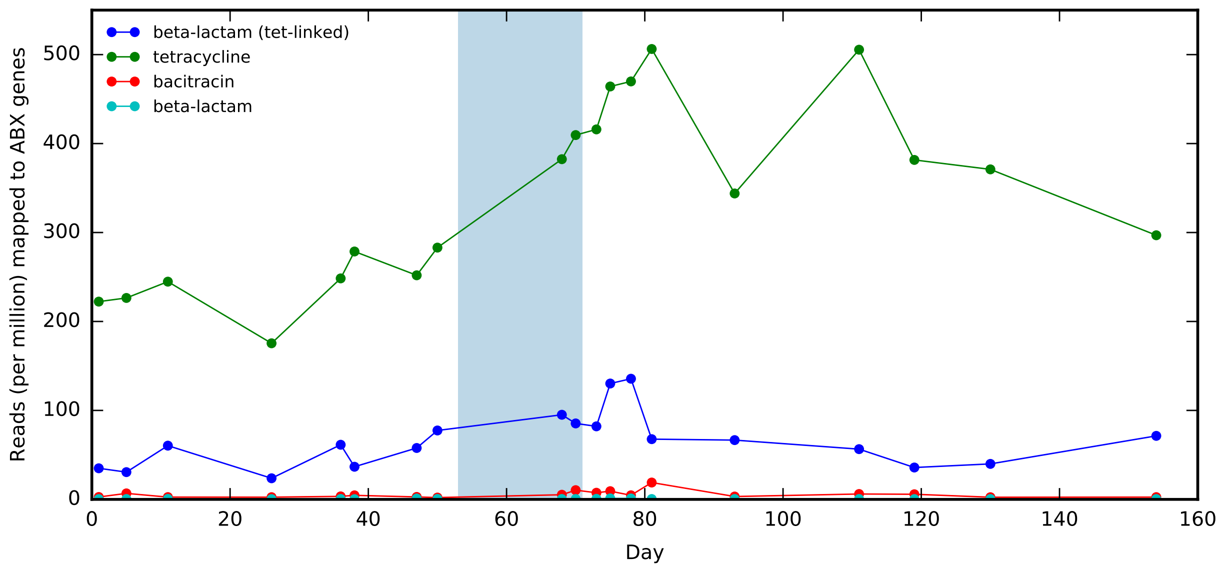

Figure S4: **Relative abundance of antibiotic resistance genes as a function of time.** Lines show fraction of reads mapping to different classes of antibiotic resistance genes in the ARGs-OAP [1] database (Methods). The shaded region indicates the antibiotic treatment period.

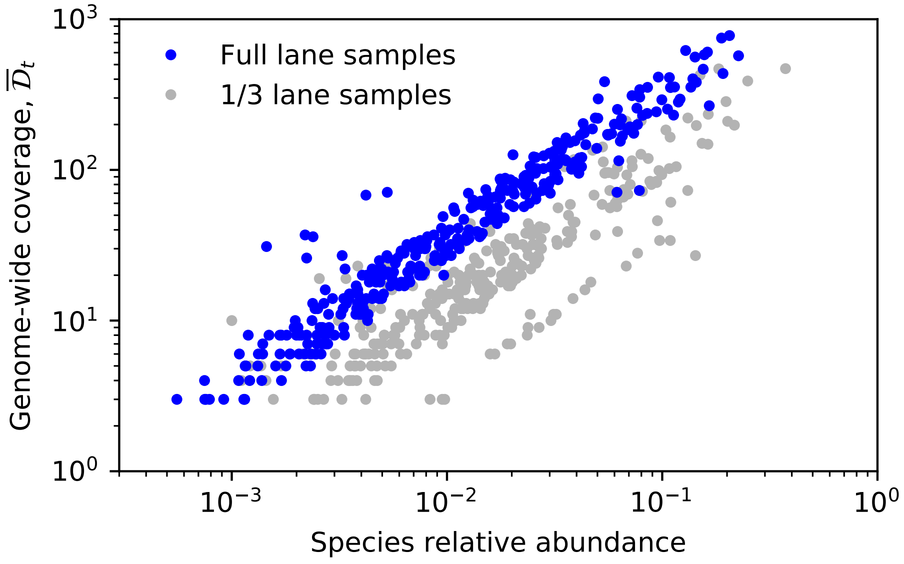

Figure S5: **Reference genome coverage as a function of species relative abundance.** Symbols denote the typical number of unique read clouds per site, $\overline{\mathcal{D}}_t$, for each reference genome in each sample as a function of the estimated relative abundance of the species in that sample. Blue points indicate samples that were sequenced to a high target coverage (Supplemental Table S1).

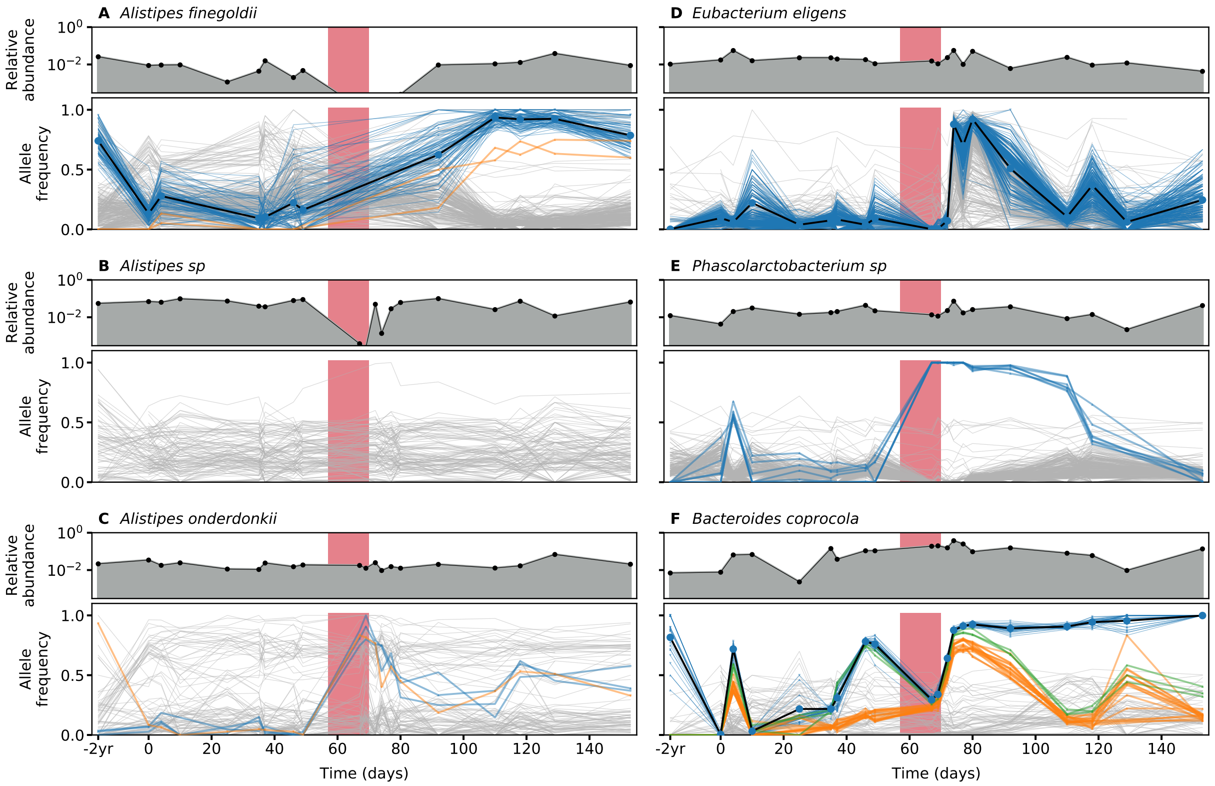

Figure S6: **Analogous version of Fig. 3 for more example species (Part 1/3).** The -2yr timepoint is estimated using data from a previous study that sampled the same individual [3].

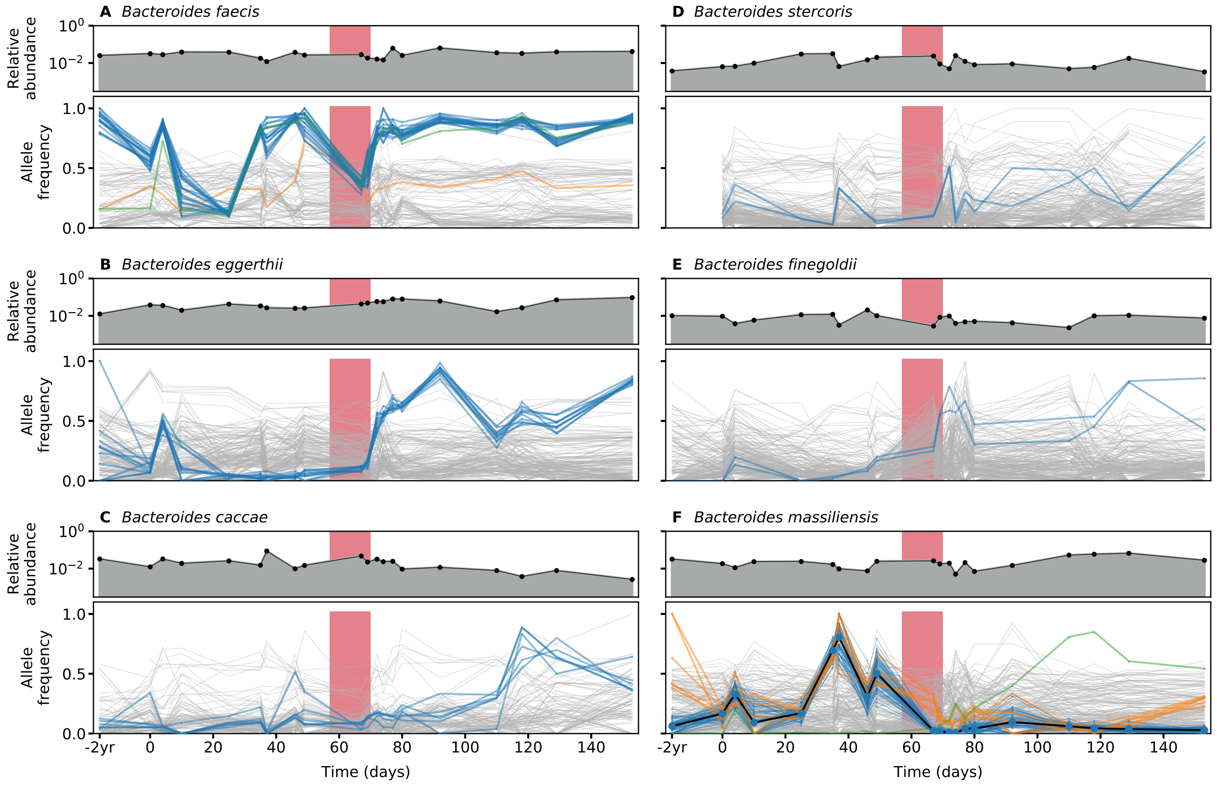

Figure S7: **Analogous version of Fig. 3 for more example species (Part 2/3).**

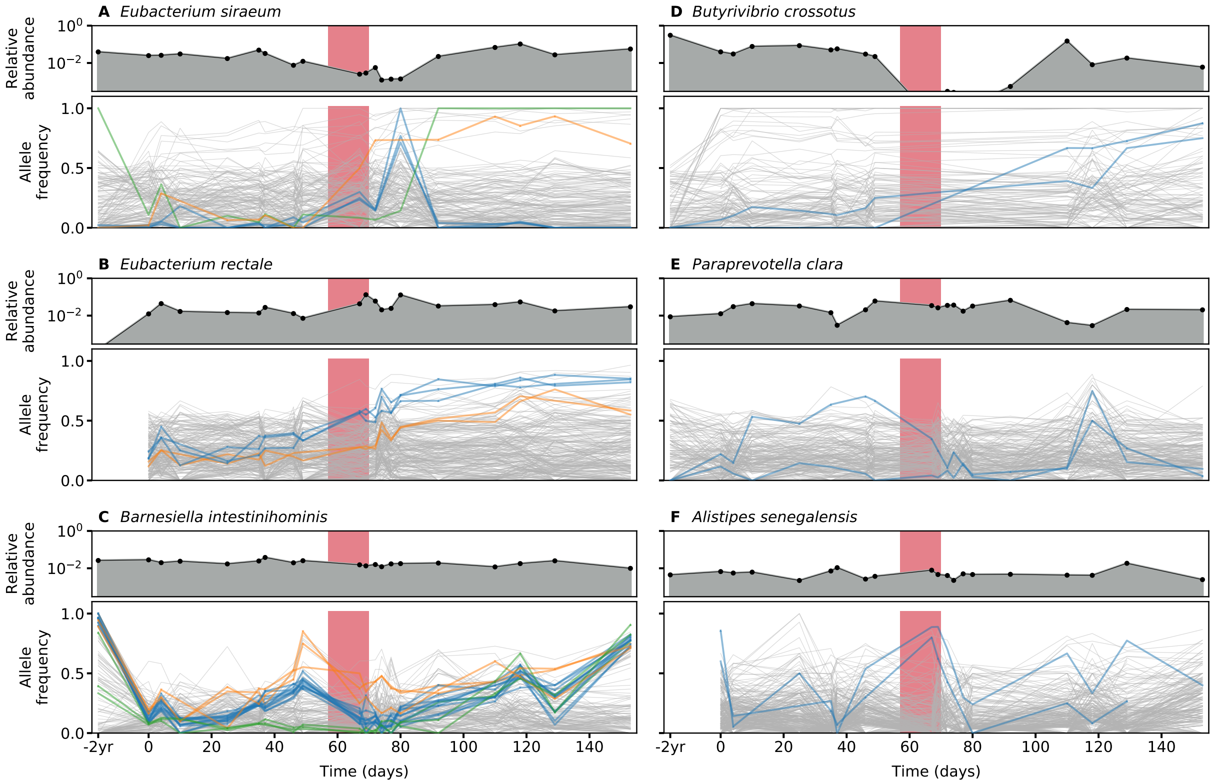

Figure S8: **Analogous version of Fig. 3 for more example species (Part 3/3).**

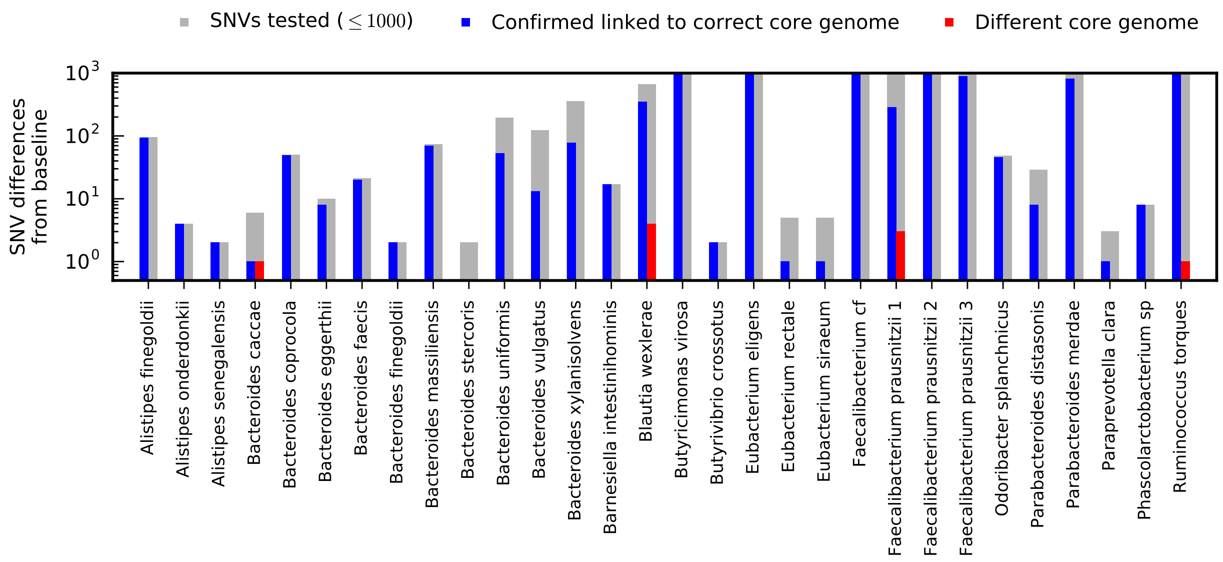

Figure S9: **Read clouds link putative within-species sweeps to the correct genomic backbone.** For each of the significant SNV differences in Fig. 2 (grey bars), we calculated levels of read cloud sharing with core genes from all of the species in our reference panel; for species with $\geq 1000$ sweeping SNVs, we chose a random subset of 1000 to analyze. For both the reference and alternate alleles of these SNVs, we retained the five genes with the largest number of shared read clouds across all timepoints (and at least 5 shared read clouds in total). Linkage to the correct species was confirmed if, for both alleles, the majority of the shared read clouds were derived from core genes in the correct species (blue). Linkage to a different species was inferred if one or both alleles had only a minority of shared read clouds from the correct species (red). Across species, we observe an average positive confirmation rate of $\approx$80% and a negative confirmation rate of <1%, with most of the negatives clustering in just a few species.

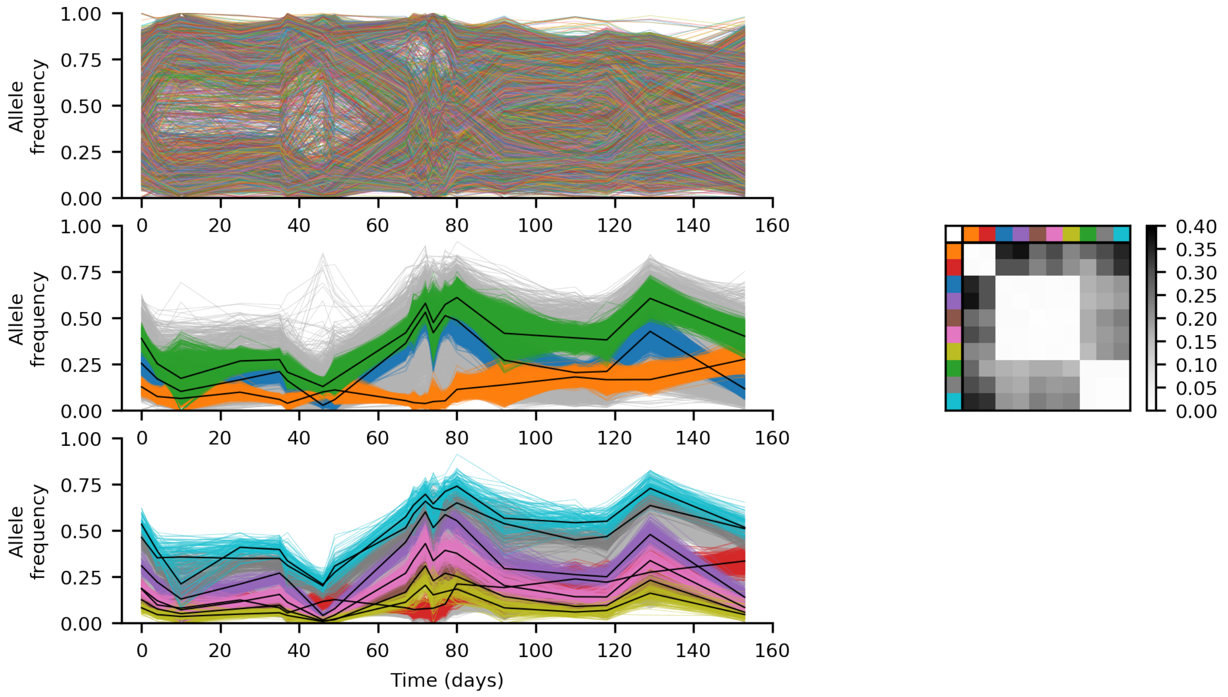

Figure S10: **Read clouds reveal haplotype structure in *B. vulgatus*.** Top panel: Allele frequency trajectories for ≈15,000 SNVs in the resident *Bacteroides vulgatus* population. Each SNV is assigned a random color for visualization purposes, and SNVs are polarized to show the frequency of the minor allele across a larger cohort of hosts in Ref [4]. Middle panel: Trajectory based clustering in Section 8 identifies three SNV clusters that contain ≈85% of all the SNVs in the top panel (colored lines). Remaining SNVs are shown in light grey for comparison. Bottom panel: Remaining SNV clusters with $\geq$ 100 SNVs per cluster (colored lines) account for ≈13% of SNVs. All other SNVs are shown in grey for comparison. (right) Fraction of read clouds supporting non-perfect linkage for SNVs in different pairs of clusters from the middle and bottom panels. This suggests that the clusters in the bottom panel are actually in perfect linkage with the major clusters in the middle panel.

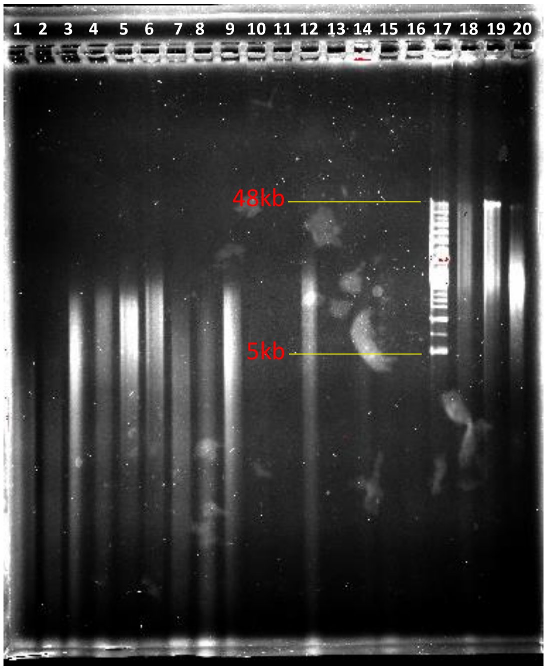

1. Human MicroBiome Project Method, dog stool sample
2. Human MicroBiome Project Method, cat stool sample
3. Human MicroBiome Project Method, human stool sample
4. MoBio PowerFecal Kit, dog stool sample
5. MoBio PowerFecal Kit, cat stool sample
6. MoBio PowerFecal Kit, human stool sample
7. MoBio PowerSoil Kit, dog stool sample
8. MoBio PowerSoil Kit, cat stool sample
9. MoBio PowerSoil Kit, human stool sample
10. QIAamp DNA stool mini Kit, dog stool sample
11. QIAamp DNA stool mini Kit, cat stool sample
12. QIAamp DNA stool mini Kit, human stool sample
13. Qiagen HMW MagAttract Kit, dog stool sample
14. Qiagen HMW MagAttract Kit, human stool sample
15. Qiagen Blood and Tissue Kit, dog stool sample
16. Qiagen Blood and Tissue Kit, human stool sample
17. CHEF 5-48kb Ladder
18. 10X Genomics standard control DNA ( DNA extracted from mammalian cells)
19. MagAttract kit control sample (DNA extracted from primate peripheral blood mononuclear cells)
20. AllPrep kit control sample (DNA extracted from primate peripheral blood mononuclear blood cell)

Figure S11: Pippin pulse gel showing the size of the HMW DNA extracted from human, dog, and cat stool samples using different extraction methods.

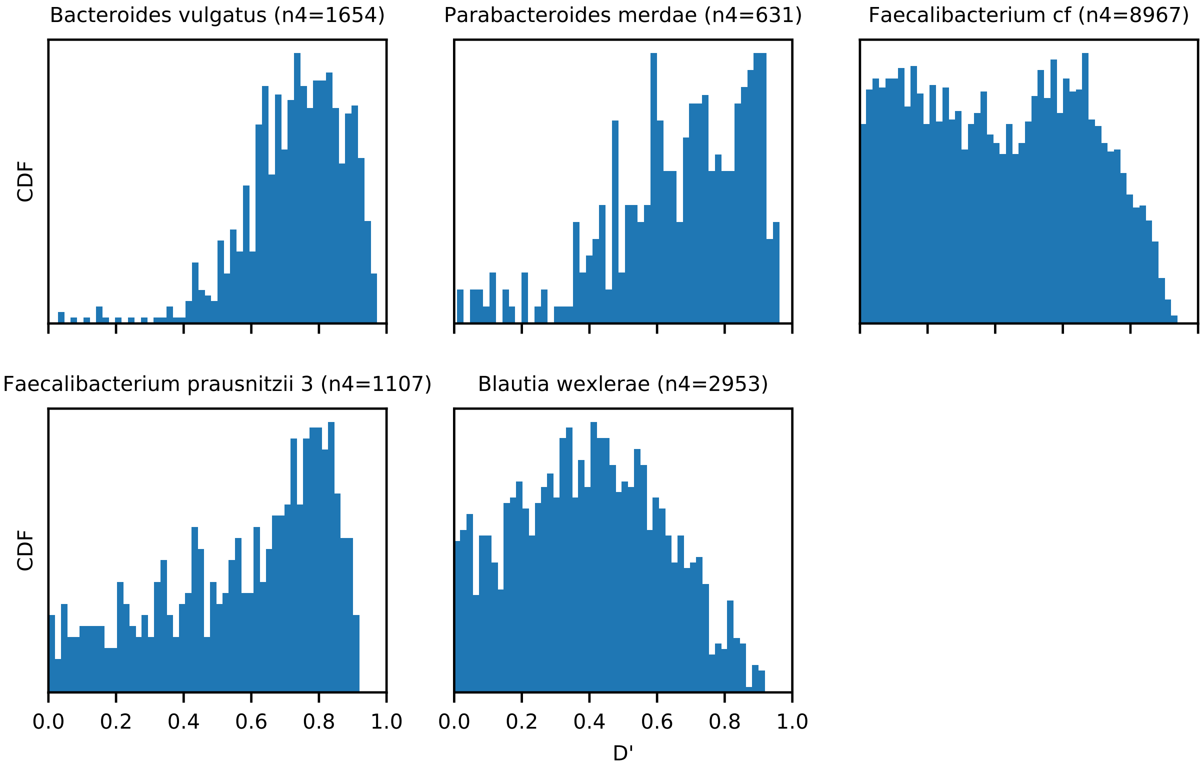

Figure S12: **Linkage disequilibrium between SNVs that fail the 4 haplotype test.** Each panel shows the distribution of $D'$ [5] for the subset of SNV pairs in Fig. 4C where all four haplotypes are observed. This measure is defined so that $D' = 1$ in the absence of recombination, and $D' \approx 0$ for unlinked loci. This distribution was calculated for all species in Fig. 4C with at least 100 4-haplotype SNV pairs. In most of these cases, violations of the 4 gamete test still cluster near $D' \approx 1$, with possible exceptions in one of the *Faecalibacterium* and *Blautia* populations.

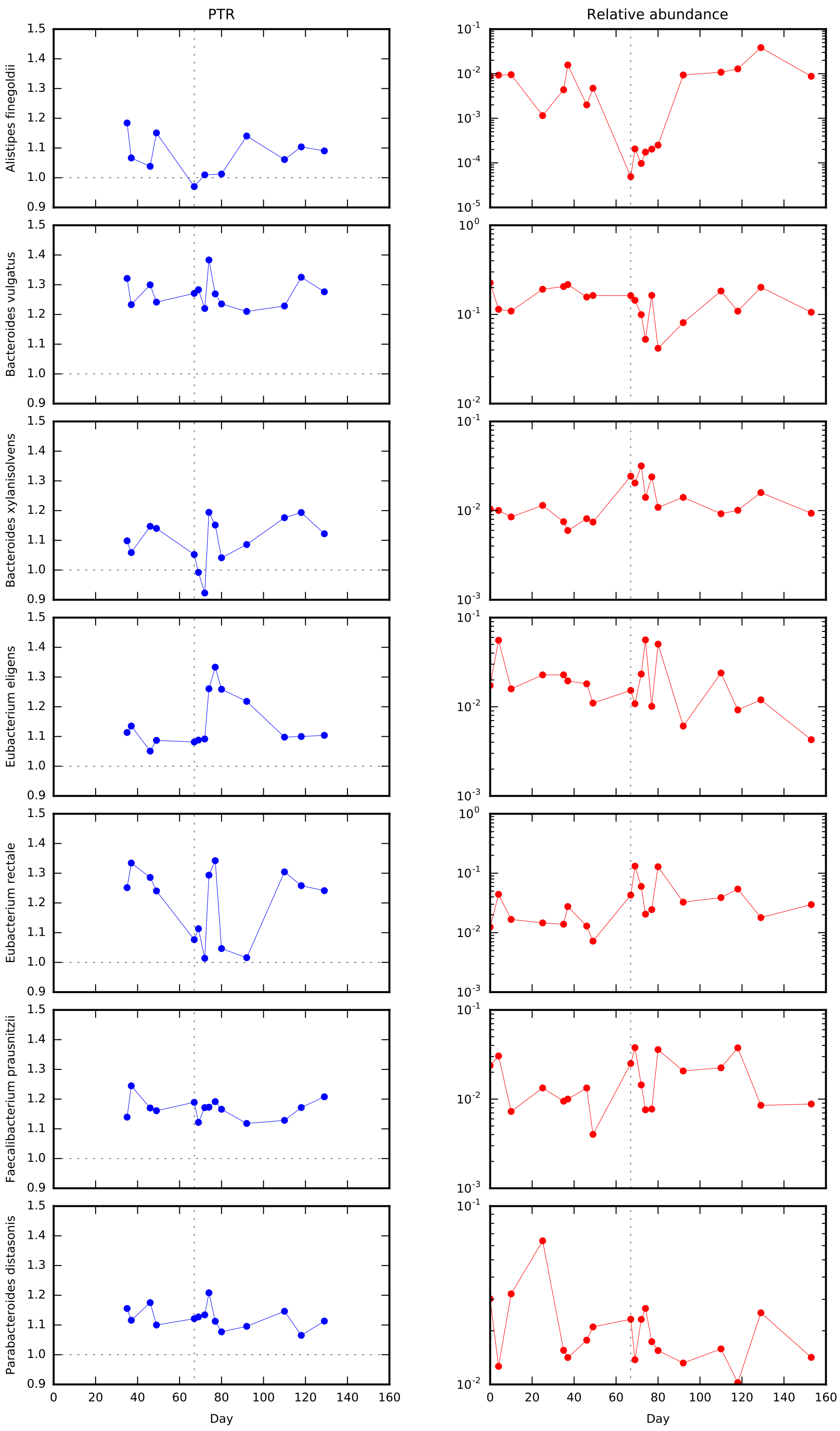

Figure S13: **Replication origin peak-to-trough ratio (PTR) and relative abundance for select species.** PTR values were obtained from the software provided by Ref [2]. Data are shown for all species that had an entries in both the Ref [2] and MIDAS databases, and had relative abundance $\geq 1\%$ in at least one timepiont. The dashed line denotes the first sample from the antibiotic treatment period.

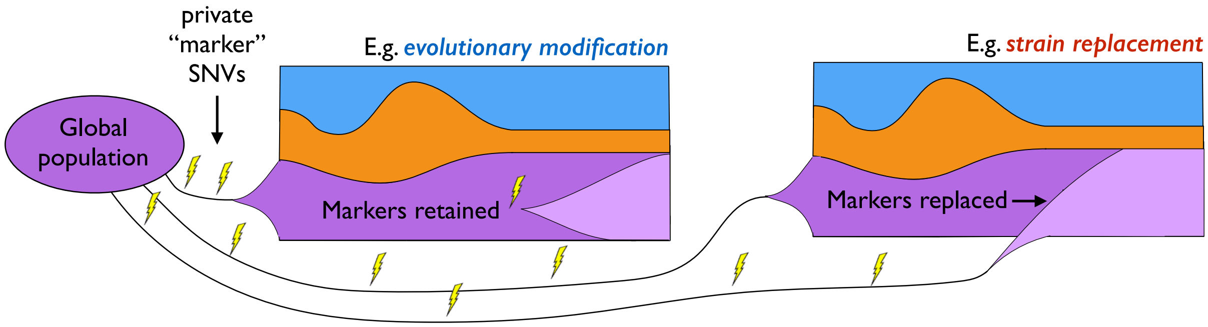

Figure S14: **Schematic illustration of private marker SNV sharing over time.** Two different within-host scenarios are depicted, which illustrate the concepts of evolutionary modification (left) and replacement by distantly related strains (right). In both cases, the within-host populations of the purple species are founded by single strains from the global population, each of which is characterized by a different set of private marker SNVs (yellow symbols). During an evolutionary modification event (left), the majority of these private marker SNVs should remain at high frequencies, while a strain replacement event (right) should cause the previous private markers SNVs to be replaced by the private markers SNVs from the invading strain. This characteristic behavior was used to motivate the private marker SNV sharing statistic defined in Section 5.5.2

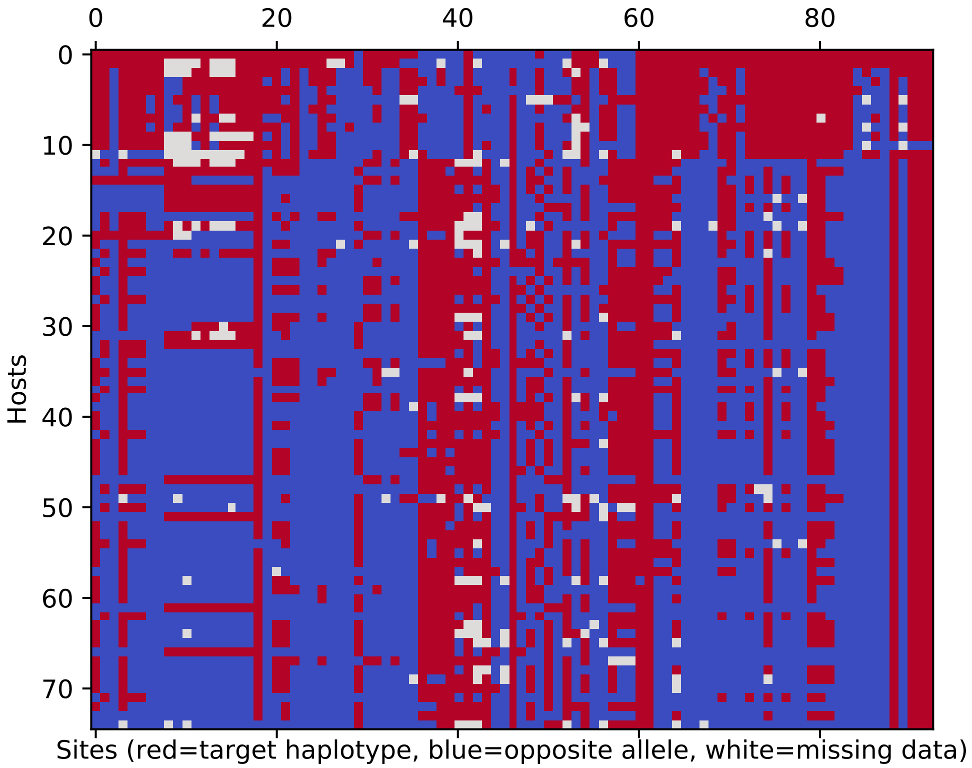

Figure S15: **Co-inheritance of *A. finegoldii* SNV differences across a larger cohort.** The SNV differences in the *A. finegoldii* example in Fig. 3A are tracked across the panel of quasi-phaseable hosts in Ref [4]. For each host/site combination, red indicates that the allele shown in Fig. 3A is also the dominant allele in that host, while blue shows the opposite; white tiles denote missing data. These data show that subsets of the SNV differences in Fig. 3A are often co-inherited together in larger haplotype blocks.

### Experimental Methods

#### 1 Study design and sample collection

Stool samples were collected every two days from a 62-year-old male over a 5-month period. Samples were collected in a 50 ml conical tube and stored on dry ice immediately after collection. Of these, 19 timepoints were selected for High Molecular Weight (HMW) DNA extraction, linked-read library preparation and sequencing (Supplemental Table S1). Approximately 200-250mg of stool samples were aliquoted and placed on dry ice in a 2ml Eppendorf tube for HMW DNA extraction.

#### 2 High molecular weight DNA extraction

High Molecular Weight (HMW) DNA was extracted from stool samples for all the study timepoints. In order to optimize the HMW DNA extraction method, seven extraction methods covering a range of methodological types were evaluated. These methods included the Human Microbiome Project extraction method [6], MoBio Powersoil DNA isolation kit, QIAamp DNA stool mini kit, Phenol Chloroform DNA isolation, Qiagen HMW DNA MagAttract, MoBio PowerFecal DNA isolation kit, and QIAgen DNeasy Blood and Tissue kit 1-3 (Supplemental Table S2). DNA extraction methods were followed according to the manufacturer's instructions. Three methods, Phenol Chloroform DNA isolation, Qiagen HMW DNA MagAttract, and QIAgen DNeasy Blood and Tissue kit, did not have homogenization steps, so a general homogenization process was adapted from Ref [7] and performed prior to these kits. In this process, 500uL of sterile PBS, 0.35 g of 0.5 mm diameter zirconia/silica beads, and 250 mg of frozen feces were added to a sterile 1.5 mL Eppendorf tube and vortexed at 3200 rpm horizontally for 4 minutes. The samples were centrifuged at 200g for 5 minutes at room temperature. 200 uL of supernatant was removed and placed in a sterile 1.5 mL Eppendorf tube. Following this step, the samples were processed according to the manufacturer's instructions. Samples were stored in 4°C to prevent shearing from freeze-thaw cycles.

***HMW DNA quality evaluation.*** Fragment Analyzer (Advanced Analytical Technologies, Inc., Ankeny, IA) was used to evaluate the size and quality of HMW DNA from each of the timepoints (Supplemental Data S5 and Supplemental Data S6). However, prior to final evaluation using Fragment Analyzer, the following steps were conducted for preliminary evaluation. Primary Extracted DNA concentrations were determined using a Broad Range assay on the Qubit Fluorometer 1.0 (Invitrogen Co., Carlsbad, CA) and purity was found using spectrophotometry at 260/280 and 260/230 absorbance ratios with Nanodrop 1000 Spectrophotometer (Thermo Fischer Scientific, Waltham, MA). This information was then used to run a pulse field gel electrophoresis with the Pippin Pulse (Sage Science Inc., Beverly, MA), providing greater separation of HMW DNA fragments. A gel was made and run with the samples according to the manufacturer's instructions for the 5-80 kb range. Each well had 100 ng of sample DNA along with Thermo Scientific 6X DNA loading dye (Thermo Fischer Scientific, Waltham, MA). One well included the CHEF 5-48kb ladder (Bio-Rad Laboratories, Pleasanton, CA) and three wells were used for genomic controls from three kits of known DNA length, 10XG control HMW DNA (10X Genomics Inc., Pleasanton, CA), MagAttract, and AllPrep. The gels were imaged with ChemiDoc MP Imaging System (Bio-Rad Laboratories, Pleasanton, CA) and visually evaluated to determine which samples and extraction methods had the longest length of DNA. Additionally, the samples were run through a High Sensitivity DNA assay using an Agilent Bioanalyzer 2100 (Agilent Technologies, Santa Clara, CA) to evaluate distribution of DNA fragments between 35 bp and 10380 bp. Samples that had greater fluorescence peaks at the 10380 bp end of the range were taken to have larger numbers of long DNA fragments than samples that had peaks throughout the range below 10380 bp. Since Bioanalyzer is not the optimal device for evaluation of extremely large fragments of DNA, other methods including Pippin Pulse Gel (Sage Science Inc.) were used to evaluate the size of microbial DNA extracted from the stool samples (Supplemental Fig. S11). QIAamp DNA stool mini Kit method (Skt 12) was used for extraction of HMW DNA from all the timepoints. An evaluation of HMW DNA size was conducted on samples from all the timepoints before 10X Genomics linked-read library preparation (Supplemental Data S5 and Supplemental Data S6).

#### 3 Linked-read library preparation and sequencing

We used 10X Genomics Chromium™ to barcode HMW fragments from 19 longitudinal samples, which were then sequenced on an Illumina HiSeq4000 platform, generating 151bp paired end reads. The target sequencing coverage for each sample is listed in Supplemental Table S1.

#### 4 Postprocessing and read cloud assignment

Following sequencing, 16bp of 10XG barcodes were removed along with sample indices. Using FastQC, we conducted quality control (QC) analysis of the sequencing reads to empirically determine whether the reads contained artifacts. We found the first 20 bp and 25 bp (R1 and R2 respectively) of each read pair had 5-mer or 7-mer sequence repeats, and these were subsequently trimmed for all reads. The 10X barcode strings were then added to the sequence identifier field in the trimmed FASTQ files so that they could be tracked in downstream steps of the pipeline.

Reads were assigned to read clouds using a lightweight error correction algorithm on the extracted barcode strings. For each sample, we created a list of all observed barcode strings and the number of reads in which they were observed. These data were then used to seed our initial set of read clouds. Iterating through barcodes in descending order, we created a new read cloud cluster if the barcode had an edit distance $>1$ from all previous read cloud entries. Barcodes with an edit distance of 1 were error corrected to the corresponding read cloud. After this initial clustering step, we excluded all read clouds where $\geq 80\%$ of the bases in the barcode label were identical, or where $\geq 50\%$ of the bases were ambiguous (`N`). Finally, we excluded all read clouds that had fewer than 10 reads. Given the observed distribution of barcode abundances, these barcodes typically account for $\leq 10\%$ of the reads in a given sample; excluding these read clouds helps to eliminate residual errors that were not caught by the initial error correction step.

### Computational Methods and Analysis

#### 5 Metagenomic pipeline

Metagenomic sequences were analyzed using a reference-based approach similar to the one we employed in Ref [4]. This pipeline utilizes the MIDAS software package [8] (software version 1.2.1, reference database version 1.2) for the initial stages of species abundance quantification, sequence alignment, and SNV detection, with several additional layers of filtering implemented in custom postprocessing scripts [4]. We further extended this pipeline to track the read cloud label (barcode) associated with each mapped short read. These steps are described in more detail below. Whenever possible, we attempted to use the same thresholds and parameter settings for this pipeline that we employed in our previous study [4], with occasional modifications to account for the additional timepoints and linked read nature of our data. All associated code is available on GitHub (`https://github.com/bgoodlab/highres_microbiome_timecourse`).

##### 5.1 Quantifying relative abundance at the species level

To estimate the relative abundances of the species in a given sample, MIDAS first maps the short sequencing reads to a panel of universal single-copy marker genes from each of the species in the MIDAS reference genome database [8]. The relative abundance of each species is then estimated from the coverage of its associated marker genes. These estimates were used for the species abundance trajectories in Figs. 1-3.

The species abundance estimates were also used to create a personalized panel of reference genomes for subsequent read mapping. Reference genomes were included for all species that had an average marker gene coverage $\geq 3$ in at least 1 timepoint. By using the same set of reference genomes across all timepoints, we aim to reduce spurious temporal variation caused by different effective mapping parameters in different samples. In our downstream analyses, we further restricted our attention to the subset of species in the reference panel with marker gene coverage $\geq 10$ in at least 3 timepoints, as we have previously shown that coverages of this magnitude are usually necessary to accurately detect SNV differences within species over time [4].

##### 5.2 Quantifying gene content for each species

We next quantified the gene content of each species in the personalized panel of reference genomes defined above. For each timepoint, MIDAS maps the short sequencing reads to a database of gene families (or *pangenome*) constructed from genes in sequenced isolates [8], using the same default parameters as Ref [4]. For each species, the average coverage was reported for each gene family and each timepoint, as well as for a panel of universal, single-copy marker genes. The copynumber $c_{gt}$ of gene $g$ at timepoint $t$ defined as the ratio between these two quantities.

We used these copynumber values to define a "core" genome for each species. We found it useful to distinguish between two different notions of a core genome. The first, which we call the ***resident genome***, contains the genes that (i) are present in the reference genome and (ii) have near single copynumber in a majority of the timepoints from the sampled host. Specifically, we required these genes to have $0.5 \leq c_{gt} \leq 2$ in more than 70% of the timepoints where the species itself had sufficient coverage (i.e., when marker gene coverage was $\geq 10$). To reduce cases of mismapping from other species in the sample, we excluded all genes whose copynumber exceeded 3 in any timepoint, and we further excluded the blacklist of putatively shared genes identified in Ref [4].

We then defined the ***core genome*** of a given species to be the subset of genes in the resident genome that are shared across most strains in other hosts. Specifically, we used the intersection between the resident genome and the set of core genes identified in our previous study [4], which were present at near single copy in $\geq 90\%$ of eligible metagenomes across a large human cohort.

#### 5.3 Identifying SNVs within species

We next identified single nucleotide variants (SNVs) within each species. We first used MIDAS to map the short sequencing reads to the personalized reference panel in Section 5.1, using the default parameters described in Ref [4]. MIDAS then reports the total sequencing coverage $D_{it}$ at each site $i$ in each timepoint $t$ [8]. Following our approach in Ref [4], we then used the distribution of coverage across the genome to obtain a measure of the typical overage at a given timepoint, $\overline{D}_t$, defined as the median of all protein coding sites with nonzero coverage. Timepoints with $\overline{D}_t < 5$ were excluded from further analysis, as were species with fewer than 4 passing timepoints. We then used the timepoint-specific estimates of $\overline{D}_t$ to refine the alignment step above, masking all sites in a given timepoint with $D_{it} < \overline{D}_t/3$ or $D_{it} > 3\overline{D}_t$. Sites that were masked in more than 30% of timepoints were excluded from further analysis. We further restricted our attention to protein coding sites in the resident genome, as defined above.

For each of the retained sites, MIDAS reports reference and alternate allele counts, $R_{it}$ and $A_{it}$, at each timepoint. In the minority of cases where multiple alternative alleles were present, these were merged into a single alternate class. To reduce bias induced by the choice of the reference genome, we re-polarized the alleles based on our previous measurements across a large human cohort [4], swapping the reference and alternate alleles so that the reference coincides with the cohort-wide consensus. Following repolarization, SNVs were retained for downstream analysis if $A_{it} \geq 0.1 D_{it}$ in at least one timepoint. (Note that this includes sites that are locally fixed in the focal host, but where the host-wide consensus is the minority allele in our larger cohort. These sites are critical for the private marker SNV tracking in Fig. 5.)

##### *5.3.1 Quantifying genetic diversity over time*

To quantify the levels of genetic diversity within each species in Fig. 2, we used a metric of intermediate-frequency polymorphism similar to the one we employed in Ref [4]. For each timepoint with $\overline{D}_t \geq 20$, we calculated the fraction of unmasked sites in the core genome with allele frequencies in the range $0.2D_{it} \leq A_{it} \leq 0.8D_{it}$. Timepoints with $\overline{D}_t < 20$ were excluded. By restricting our attention to intermediate frequency polymorphisms, we aim reduce the contribution of sequencing and mapping errors that would ordinarily limit resolution to $\gtrsim 10^{-3}$ SNVs/bp.

##### *5.3.2 Synonymous and nonsynonymous variants*

To infer the protein coding effect of each SNV, we used MIDAS's annotation of 1-fold, 2-fold, 3-fold, and 4-fold degenerate sites, which are inferred from the codon reading frame of the annotated genes in each reference genome [8]. These annotations are shown for the temporally variable SNVs in Fig. 5. Our definition of nonsynonymous SNVs includes all SNVs that change the amino acid sequence of the protein, while synonymous SNVs leave the amino acid sequence unchanged. Under this definition, 1-fold degenerate sites are always nonsynonymous, while 4-fold degenerate sites are always synonymous. The protein coding effect of 2- and 3-fold degenerate sites can be either synonymous or nonsynonymous depending on the specific alleles involved. To avoid ambiguity, we primarily focused on comparisons of 1-fold and 4-fold degenerate sites.

We used these annotations to calculate an effective $dN/dS$ value for the temporally varying SNVs identified in Fig. 3. To do so, we divided the observed ratio of 1D and 4D SNVs for a given species by the median ratio of unmasked 1D and 4D sites in the resident genome of that species across each timepoint. The $dN/dS$ estimates for the six example species are listed in Supplemental Data S1.

###### 5.3.3 Allele prevalence across hosts

To estimate the prevalence of alleles within the broader human population, we used the allele prevalence values obtained from our previous study of a large human cohort [4]. In that work, prevalence was defined as the fraction of metagenomic samples in which the allele attained majority frequency. The prevalence values in Fig. 5 were polarized to be consistent with the allele frequency trajectories in Fig. 3 (i.e., prevalence refers to the allele displayed in the frequency trajectory).

##### 5.4 Tracking read clouds associated with SNVs and genes

To leverage the additional linkage information encoded in the read cloud labels, we developed a new MIDAS module to track read cloud labels stored in the read name field in the underlying FASTQ and BAM files. Read cloud labels were recorded at both the individual gene and SNV levels. For each timepoint, we recorded the read cloud labels for each read that mapped to one of the gene families in Section 5.2 with MAPQ$\geq$ 20. (The additional MAPQ filter is necessary to prevent ambiguously mapping reads from being mapped to a different location than the other reads in the same read cloud.)

To efficiently track read cloud labels at the SNV level, we focused on a subset of target SNVs from Section 5.3 above that (i) had $A_{it} \geq 0.1 D_{it}$ in at least 3 timepoints, and (ii) were unmasked in at least 10 timepoints. For each of these SNVs, we separately recorded the read cloud labels for the reads that mapped to the reference and alternate allele in each timepoint, as well as the total number of matching reads for each read cloud.

In a small fraction of cases, both alleles could be observed in different reads from the same read cloud. Although this could in principle arise from read cloud impurities (see Section 7 below), a more likely explanation is that the library preparation step generated multiple short reads from the same input molecule (e.g. during the PCR step), and then sequencing or PCR errors generated reads supporting the opposite allele. We used this intuition to implement a simple form of error correction: for read clouds that contained a mixture of both alleles at a given site, we only assigned the read cloud to the allele with the largest number of supporting reads. The corrected read clouds were also recorded so that they could later be used to estimate the rates of sequencing or PCR errors in a site-specific manner. In particular, for each SNV $i$, we estimated an error rate

$$p_{\mathrm{err},i} = \frac{n_{\mathrm{err}}}{n_{D\geq 2}} \,, \tag{1}$$

where $n_{\mathrm{err}}$ is the total number of error corrected read clouds at site $i$ across all timepoints, and $n_{D\geq 2}$ is the total number of read clouds at site $i$ that have at least two reads covering that site. Average values of $p_{\mathrm{err},i}$ are on the order of $10^{-3}$, but a small fraction of sites can reach much higher values.

#### 5.5 Quantifying frequency trajectories of SNVs

We estimated the allele frequencies of SNVs using the read cloud labels recorded in Section 5.4. For a given site $i$ and timepoint $t$, let $\mathcal{A}_{it}$ and $\mathcal{R}_{it}$ denote the total number of unique read cloud labels that map to the alternate and reference alleles, respectively. (Note the cursive font, which distinguishes these quantities from the read-based counts $A_{it}$ and $R_{it}$ defined in Section 5.3.) An analogous version of the total coverage can then be defined as $\mathcal{D}_{it} \equiv \mathcal{A}_{it} + \mathcal{R}_{it}$. While the allele counts derived from reads and read clouds are expected to be correlated with each other, the read-based version will also contain an additional source of noise arising from the PCR amplification step in the library creation protocol. We therefore sought to use the read cloud versions of the allele counts as much as possible, since these are expected to more closely approximate the idealized sampling of individual molecules in our original pool of DNA.

We therefore repeated several of the filtering steps defined for the read-based counts in Section 5.3 above. We formed a read-cloud-based estimate of the typical coverage at a given timepoint, $\overline{\mathcal{D}}_t$, defined as the median value of $\mathcal{D}_{it}$ across all target SNVs. Timepoints with $\overline{\mathcal{D}}_{it} < 5$ were excluded from further analysis. As above, we then masked individual sites in a timepoint if $\mathcal{D}_{it} < \overline{\mathcal{D}}_t/3$ or $\mathcal{D}_{it} > 3\overline{\mathcal{D}}_t$, and SNVs that were masked in more than 30% of timepoints were excluded from further analysis. Based on these data, we estimated the allele frequency trajectory for each SNV using the simple plug-in estimator,

$$\hat{f}_{it} = \mathcal{A}_{it}/\mathcal{D}_{it} \, . \tag{2}$$

These frequency trajectories are shown in Fig. 3, and are used in various additional analyses described below.

##### 5.5.1 *Detecting SNV differences over time*

To quantify shifts in the genetic composition of individual species, we searched for SNVs that underwent large changes in allele frequency between different timepoints; this indicates a full or partial "sweep" of the allele through the resident population of interest. Given our study design, we were particularly interested in cases where this "sweep" occurred between one of the baseline timepoints and a later stage of the study (e.g. antibiotic treatment). The primary difficulty lies in disentangling these true allele frequency changes from spurious signals produced by the finite sampling of sequencing reads in each timepoint. The uncertainty in the sample frequency scales as $\sigma_f \sim \mathcal{D}^{-1/2}$, which grows increasingly small at higher coverages. However, the effective error rate can be amplified by the large number of sites and temporal comparisons that we must search through. To overcome this issue, we used an extension of the approach we developed in Ref [4] for detecting genetic differences between pairs of timepoints.

For each pair of timepoints $t_1 < t_2$, we searched for SNVs where the minor allele transitioned from less than 20% frequency in the first timepoint to more than 70% in the second timepoint. In terms of the allele counts, this requires either (i) $\mathcal{A}_{it_1} \leq 0.2\mathcal{D}_{it_1}$ and $\mathcal{A}_{it_2} \geq 0.7\mathcal{D}_{it_2}$, or (ii) $\mathcal{A}_{it_1} \geq 0.8\mathcal{D}_{it_1}$ and $\mathcal{A}_{it_2} \leq 0.3\mathcal{D}_{it_2}$. Under the null hypothesis of no allele frequency change (i.e., the true frequency is the same at both timepoints), the probability of observing such an event by chance through sampling noise is given by

$$\begin{aligned} P_i(t_1,t_2) = & \underbrace{F(0.2\mathcal{D}_{it_1}|\mathcal{D}_{it_1},\overline{f})F(0.3\mathcal{D}_{it_2}|\mathcal{D}_{it_2},1-\overline{f})}_{\text{from } \leq 0.2 \text{ to } \geq 0.7} \\ & + \underbrace{F(0.2\mathcal{D}_{it_1}|\mathcal{D}_{it_1},1-\overline{f})F(0.3\mathcal{D}_{it_2}|\mathcal{D}_{it_2},\overline{f})}_{\text{from } \geq 0.8 \text{ to } \leq 0.3} \, , \end{aligned} \tag{3}$$

where $\overline{f} = (\mathcal{A}_{it_1} + \mathcal{A}_{it_2})/(\mathcal{D}_{it_1} + \mathcal{D}_{it_2})$ is the average allele frequency between the two timepoints and $F(k|n, p)$ is the cumulative distribution function for a binomial distribution with sample size $n$ and success probability $p$. The expected number of differences under the null hypothesis is therefore given by

$$n_{\rm err}(t_1, t_2) = \sum_i P_i(t_1, t_2)\,, \tag{4}$$

which can be compared to the corresponding observed value, $n_{\rm obs}(t_1, t_2)$.

To maximize statistical power, we excluded sites from the sum in both the observed and expected counts if they met any of the following criteria

1. **High false positive rate:** $P_i(t_1, t_2) > 10^{-3}$.
2. **Insufficiently polymorphic:** the minor allele frequency is $\leq$0.2 in all timepoints.
3. **Variable copynumber:** the ratio between $\mathcal{D}_{it}$ and the typical genome-wide coverage $\overline{\mathcal{D}}_t$ differed by more than a factor of 2 between $t_1$ and $t_2$.

The last condition is useful for reducing false positives caused by mis-mapped reads from other species (see Ref [4] and Section 7.5 below). Technically speaking, the polymorphism condition has the potential to shift the null distribution away from the simple binomial form in Eq. (3), since it depends on the properties of the allele counts beyond the mean $\overline{f}$. We neglect this influence here, since the polymorphism condition can often be satisfied at sites other than $t_1$ and $t_2$.

Based on these definitions, we calculated the observed and expected SNV differences for all pairs of timepoints where $t_1$ is one of the 4 baseline samples and $t_2$ is one of the 15 remaining samples. We considered there to be sufficient evidence for SNV differences between a pair of timepoints if (i) the estimated false discovery rate, $n_{\rm err}(t_1, t_2)/n_{\rm obs}(t_1, t_2)$, is less 0.1 and (ii) the Bonferonni-corrected $P$-value for the total number of changes,

$$P \approx (4 \cdot 15) \sum_{k=n_{\rm obs}}^{\infty} \frac{(n_{\rm err})^k}{k!} e^{-n_{\rm err}}\,, \tag{5}$$

is less than 0.05. If these conditions were met, we recorded a significant SNV difference for all the SNVs that were observed to change between these two timepoints. In this way, we are effectively leveraging correlations in the dynamics of SNV trajectories across the genome to overcome some of the uncertainty involved in any single SNV trajectory.

We performed this procedure for all species that had at least 6 unmasked timepoints (i.e. with $\overline{\mathcal{D}}_t \geq 5$) and at least one baseline timepoint with $\overline{\mathcal{D}}_t \geq 10$. A total of 36 species met these criteria (see Fig. 2). For each species, the total set of temporally variable SNVs (or ***SNV differences***) was defined as the union of the SNVs identified in any significant timepoint pair. This list was used to determine the colored trajectories in Fig. 3 and the total number of SNV differences reported in Fig. 2C. We note that this method of detecting temporally variable SNVs may miss many true allele frequency changes that do not reach our stringent thresholds (e.g. the *Bacteroides vulgatus* example in Supplemental Fig. S10). The estimates in Figs. 2 and 3 should therefore be viewed as a lower bound, highlighting only the most dramatic allele frequency shifts. In principle, it should be possible to increase our detection sensitivity significantly by leveraging the correlations in the allele frequency changes across multiple timepoints, similar to Refs [9–11]. Such techniques have previously been employed only in relatively simple metagenomes derived from laboratory evolution experiments [9–11]. Extending these approaches to the more complex strain mixtures encountered in natural populations remains an important avenue for future work.

##### *5.5.2 Private marker SNVs*

To help distinguish instances of strain replacement from the evolutionary modification of resident strains, we examined the dynamics of so-called ***private marker SNVs*** across the sampled timecourse. Similar approaches have been used to quantify sharing or transmission of bacterial strains across different hosts [8, 12–14]. Here, we are effectively using the same idea to quantify transmission of bacterial strains between successive timepoints from the same host, as illustrated in Supplemental Fig. S14.

We used a generalization of the approach we developed in Ref [4] for analyzing pairs of timepoints. This approach is motivated by three basic assumptions:

1. Most of the private SNVs that are found at high frequencies at the initial timepoint are passenger mutations that were acquired by the resident strain before it colonized the host, and hitchhiked to high within-host frequencies during the colonization process. We refer to these SNVs as ***private marker SNVs***. (Some of these private SNVs may also reflect de novo mutations that have swept to high frequency in the time since colonization; we assume that these are in the minority due to the different timescales between within- and between-host evolution.)

2. There is no strong selection pressure to revert private marker SNVs through evolutionary modification (i.e., most of the alternate alleles are not strongly deleterious).

3. Replacement events draw from a sufficiently diverse pool that the new strain is unlikely to share the same private marker SNVs as the resident strain.

Together, these assumptions imply that the preservation or disruption of private marker SNVs at later timepoints can distinguish between strain replacement and evolutionary modification.

Based on this intuition, we defined the set of ***disrupted private marker SNVs*** to be the subset of temporally variable SNVs whose prevalence (defined Section 5.3.3) is exactly 1 (i.e., where the *reference* allele as shown in Fig. 3 is private). Conversely, we defined the set of ***preserved private marker SNVs*** to be the subset of private SNVs that were in the majority at all timepoints, and had a frequency $\geq$0.8 in at least 80% of timepoints. These numbers were used to construct the pie charts in Fig. 5.

### 6 Antibiotic resistance gene profiling

Since antibiotic resistance genes may have poor coverage within the default MIDAS database, we used a separate approach to calculate the levels of antibiotic resistance genes in Supplemental Fig. S4. Our approach is essentially a lightweight wrapper around the ARGs-OAP pipeline [1], which was specifically developed to detect antibiotic resistance genes in metagenomic data. Briefly, short sequencing reads were mapped against a database of antibiotic resistance genes using UBLAST [15], and putative matches refined with BLAST [16]. Reads that matched one of the genes in the database were extracted and labelled with the class of antibiotics that the gene is known to confer resistance to (e.g. tetracycline) as well as the gene family of the antibiotic resistance gene (e.g. tetQ). These data are plotted in Supplemental Fig. S4, which shows the number of reads (per million mapped) that are assigned to genes in several antibiotic classes.

### 7 Inferring genetic linkage from shared read clouds

In principle, read clouds generated from a single DNA molecule could provide direct evidence of genetic linkage over much longer distances than a typical short sequencing read. In practice, this problem is made more difficult due to the presence of *read cloud impurities*, which occur when multiple DNA molecules are labelled with the same barcode. Below, we describe our efforts to quantify the extent of read cloud impurity in our data. We then present a statistical approach for inferring genetic linkage between variants based on elevated rates of read cloud sharing across many individual read clouds.

To fix notation in the sections below, we will let $\mathscr{B}_t$ denote the total collection of read clouds that were sequenced in timepoint $t$, with $\mu \in \mathscr{B}_t$ denoting a particular read cloud from this set. For each read cloud $\mu$, we will let $D_\mu$ denote the total number of read pairs that were sequenced from that read cloud. A central challenge for our analysis is that the observed values of $D_\mu$ span several orders of magnitude. These large differences reflect both the intrinsic variation between read clouds from the same timepoint, as well as systematic differences between timepoints that were sequenced at different overall depths (Fig. 1). This has important consequences for read cloud impurity, since $D_\mu$ places strong constraints on the *realized* level of impurity in a given read cloud. For example, a read cloud with $D_\mu = 2$ can contribute reads from at most 2 fragments, regardless of the number of DNA molecules in the emulsified droplet. Since the empirical distribution of $D_\mu$ is skewed towards these low values, simple averages over $\mu$ can severely underestimate the risks of read cloud impurity when inferring genetic linkage. The methods below are designed to overcome this problem, by conditioning on the actual values of $D_\mu$ that are observed in a given situation.

#### 7.1 Empirical estimates of read cloud impurity

In metagenomic settings, crude estimates of read cloud impurity can be obtained by counting the number of species that contribute reads to a given read cloud. This is effectively a lower bound on the number of DNA fragments, which becomes increasingly tight for large and uniformly distributed communities. To calculate the effective number of species in Fig. 1, we used the species assignments implicit in the gene-level mapping in Section 5.4. For each read cloud $\mu \in \mathcal{B}_t$, we calculated the total number of reads ($r_{\mu s}$) that map gene families in species $s$ in the reference panel. If all species contribute the same number of reads, then the number of species is given by the naive estimator,

$$S_\mu = \sum_s \theta(r_{\mu s}) \,, \tag{6}$$

where $\theta(z)$ is the Heaviside step function. In practice, we opted to use the root mean squared estimator,

$$S_\mu \equiv \sqrt{\frac{\sum_s \theta(r_{\mu s})}{\sum_s \left( \frac{r_{\mu s}}{\sum_{s'} r_{\mu s'}} \right)^2}} \,, \tag{7}$$

since we expect this to be more robust in the presence of a small number of misassigned reads.

The curves in Fig. 1E were obtained by binning reads according to their total coverage $D_\mu$, and plotting the median value of $S_\mu$ for each bin. As expected, the effective number of species increases with the total coverage of each read cloud, reflecting higher realized rates of read cloud impurity with increasing $D_\mu$. The typical values of $S_\mu$ range from 2 to 20, consistent with previous work by Danko et al. [17]. These values indicate that read cloud impurity is not a rare error mode, but is instead a typical property of our read cloud data.

##### 7.2 Empirical estimates of long-range linkage within read clouds

Despite the high rates of read cloud impurity in Fig. 1, these data still contain many true examples of long range linkage. To quantify this signal, we calculated the fraction of shared read clouds between pairs of SNVs as a function of the genome coordinate distance $\ell$ between them (Fig. 4B). High rates of sharing are always expected for short distances (when both SNVs are present on the same short sequencing read). Elevated rates of sharing at intermediate distances indicate true examples of long-range linkage, and they allow us to estimate the typical length scales accessed by our HMW DNA extraction protocol.

We estimated $\ell$ using the locations of the SNVs on the reference genome. We only considered pairs of SNVs on the same contig, we estimated $\ell$ as the minimum of $\ell$ and $\ell_c - \ell$ (where $\ell_c$ is the length of the contig) to account for circular chromosomes. Although differences in synteny may cause the true value of $\ell$ to vary between strains, we expect this to have only a minor impact on the genome-wide signal in Fig. 4B. We binned SNV pairs in logarithmic intervals of $\ell$, and we calculated the sharing fraction

$$P(\ell) = \frac{\sum_{ij} \mathcal{B}_{ij}}{\sum_{ij} \mathcal{B}_i}\,, \tag{8}$$

for each bin, where $\mathcal{B}_{ij}$ is the total number of shared read clouds between SNVs $i$ and $j$ across all timepoints, and $\mathcal{B}_i$ is the total number of read clouds that cover SNV $i$ (equivalent to $\mathcal{D}_i$ in section Section 5.5). To reduce potential mapping artifacts, we excluded SNVs with $\mathcal{B}_i \leq 10$, or which fell outside the middle 80% of the distribution of $\mathcal{B}_i$ for that species.

As expected, the curves in Fig. 4B show high rates of read cloud sharing for SNVs within a typical read length of each other (∼150bp), and a second peak at ∼300bp expected from paired end reads. Beyond this point, we observe a broad shoulder that extends out to $\ell$∼8kb before rapidly decaying. This represents the additional linkage information captured by the read cloud protocol, and indicates the typical length scale ($\lesssim$ 10kb) where we can hope to observe this linkage information. This length scale is consistent with independent measurements of the fragment sizes produced by our HMW DNA extraction protocol (Supplemental Fig. S11). However, we note that only ∼10% of read clouds contain this long-range linkage information. Thus, a site must be covered by at least 10 distinct read clouds before we expect to observe any linkage beyond 100bp, and much larger coverages are required to reliably observe multiple instances of read cloud sharing. For a given total depth of sequencing, this sets a minimum abundance threshold for species where we expect read cloud sequencing to be useful.

##### 7.3 Statistical null model of read cloud sharing

The strong *global* signal in Fig. 4B, combined with the high levels of read cloud impurity in Fig. 1, has important implications for inferring local genetic linkage between specific pairs of sequences. Since typical read clouds contain several fragments, individual examples of read cloud sharing do not provide conclusive evidence for genetic linkage on their own. We must instead adopt a statistical approach, scanning for sequences with higher rates of barcode sharing than expected by chance (e.g., given the background rates of read cloud sharing in Fig. 4B).

To determine these thresholds, it is necessary to model the patterns of read cloud sharing that could be produced by read cloud impurities alone. Previous approaches have focused on the special case where the sequences are derived from pure cultures of long, diploid genomes (e.g. humans). In this section, we present an extension of this approach that can account for the highly skewed abundances of different genomes in a complex metagenomic sample.

We begin by considering some genomic feature $i$, e.g., a particular site from the reference genome panel in Section 5.1. We will refer to this as the ***focal site***. Let $\mathscr{B}_i \subset \mathscr{B}_t$ denote the total set of read clouds in which at least one read maps to the the target site, so that $\mathcal{B}_i = |\mathscr{B}_i|$ denote the total number of such read clouds. We want to formulate a null model for the number of read clouds $B_{ij} \leq B_i$ that are shared with some other ***target site*** $j$. Under the null hypothesis, the two SNVs cannot be present on the same fragment, so the only way that they can share a read cloud is if separate fragments containing these SNVs appear in the same read cloud by chance. We wish calculate the null distribution of $B_{ij}$ in this scenario, conditioned on the observed distribution of read coverages in each cloud, $\mathscr{D}_i = \{D_\mu : \mu \in \mathscr{B}_i\}$.

To do so, it is useful to first consider a random read cloud with a particular value of $D_\mu$. We assume that there are constants $p_i$ and $p_j$, and a function $F(D)$, such that the probability that $i$ and $j$ are covered by reads in $\mu$ can be written in the Poisson-like form

$$\begin{aligned} \Pr[i \in \mu | D_\mu] &\approx p_i F(D_\mu)\,, \\ \Pr[i \in \mu, j \in \mu | D_\mu] &\approx p_i p_j F(D_\mu)^2\,, \end{aligned} \tag{9}$$

The function $F(D)$ will depend on the details of the barcoding and sequencing reaction, but is crucially independent of $i$ and $j$. For example, in a simple model where every read cloud contains $\overline{M}$ fragments of size $\ell_f$, the function $F(D)$ can be written in the form

$$F(D) = \overline{M}\left(1 - e^{-D\ell_r/\overline{M}\ell_f}\right) \approx \begin{cases} D\ell_r/\ell_f & \text{if } D \ll \overline{M}\ell_f/\ell_r\,, \\ \overline{M} & \text{if } D \gg \overline{M}\ell_f/\ell_r\,. \end{cases} \tag{10}$$

This suggests that Eq. (9) will be a good approximation provided that $\overline{M} \gg 1$ and $p_i\overline{M} \ll 1$. In practice, the actual shape of $F(D)$ is likely to be more complicated than the simple version in Eq. (10). We therefore treated it as an arbitrary function to be estimated self-consistently from the data.

Assuming that read clouds are generated approximately independently, we have

$$\mathcal{B}_i \sim \text{Poisson}\left(p_i \sum_{\mu \in \mathscr{B}_t} F(D_\mu)\right)\,. \tag{11}$$

This suggests that we can estimate $p_i$ from the observed values of $\mathcal{B}_i$ via

$$\hat{p}_i \approx \frac{\mathcal{B}_i}{\sum_{\mu \in \mathscr{B}_t} F(D_\mu)} \equiv \frac{q_i}{\langle F(D) \rangle}\,, \tag{12}$$

where we have defined

$$q_i \equiv \frac{\mathcal{B}_i}{|\mathscr{B}_t|}\,, \tag{13}$$

and

$$\langle F(D_\mu) \rangle_{\mu \in \mathscr{B}_t} \equiv \frac{1}{|\mathscr{B}_t|} \sum_{\mu \in \mathscr{B}_t} F(D_\mu)\,. \tag{14}$$

The conditional distribution of $\mathcal{B}_{ij}$ is therefore approximately given by

$$\mathcal{B}_{ij} \sim \text{Poisson}\left(q_j \sum_{\mu \in \mathscr{B}_i} G(D_\mu)\right)\,, \tag{15}$$

where we have defined a standardized version of $F(D)$,

$$G(D) = \frac{F(D)}{\langle F(D) \rangle} \,. \tag{16}$$

**Estimating $G(D)$.** We now describe an empirical method for estimating $G(D)$ directly from the data. To do so, we'll make use of the fact that the vast majority of SNV pairs in different species satisfy the null model above (i.e., they should never be found on the same fragment). Thus, we can try to estimate $G(D)$ by taking averages over all barcodes $\mu$. It will be helpful to introduce some new notation. Let $s(i)$ denote the species corresponding to SNV $i$, and we'll let $j \in s(i)$ and $j \notin s(i)$ indicate that SNV $j$ is or is not in the same species, respectively. For a given barcode $\mu$ (with coverage $D_\mu$), we can look at all the SNVs $i \in \mu$ that are covered by at least one read in that barcode. We'll let $n_{\mu,s}$ denote the number of SNVs of this sort that belong to species $s$. We'll also let $Q_s = \sum_{i \in s} q_i$ and $Q = \sum_s Q_s$.

With these definitions, for each SNV $i$ in a barcode $\mu$, the observed number of shared barcodes in other species is given by $n_{i,other} = n - n_{s(i)}$. Furthermore, its expectation is given by

$$E[n_{i,other} | i \in \mu, D_\mu] = (Q - Q_{s(i)}) G(D_\mu) \,. \tag{17}$$

Thus, for all $\mu$ with $D_\mu \approx D$, we can estimate

$$\hat{G}(D) = \frac{\sum_{\mu: D_\mu \approx D} \sum_{i \in \mu} \frac{n - n_{s(i)}}{Q - Q_{s(i)}}}{\sum_{\mu: D_\mu \approx D} \sum_{i \in \mu} 1} \,, \tag{18}$$

or after coarse-graining at the species level,

$$\hat{G}(D) = \frac{\sum_{\mu: D_\mu \approx D} \sum_{s \in \mu} n_s \cdot \frac{n - n_s}{Q - Q_s}}{\sum_{\mu: D_\mu \approx D} \sum_{s \in \mu} n_s} \,. \tag{19}$$

We used this formula to estimate $G(D)$ separately for each timepoint $t$, binning read clouds according to $D_\mu$ using 16 equally spaced logarithmic bins from $D = 10$ to $D = 10^4$. For bins with less than $10^4$ read clouds, we replaced Eq. (19) with the value of $\hat{G}(D)$ calculated from the nearest bin that exceeded this occupation threshold. The estimated values of $\hat{G}(D)$ are shown in Supplemental Fig. S16 below. As anticipated from our discussion above, the estimated values of $\hat{G}(D)$ can vary dramatically, both when comparing different timepoints and within timepoints when comparing read clouds with different values of $D_\mu$.

#### 7.4 Inferring linkage between pairs of SNVs

Using the null model above, we developed a statistical approach for quantifying genetic linkage between pairs of SNVs. We first searched for SNV pairs with significantly elevated rates of read cloud sharing, regardless of the allelic combinations involved. We then examined read cloud sharing at the allelic level for this subset of significantly linked SNV pairs.

##### *7.4.1 Quantifying linkage at the SNV level*

For each pair of SNVs $i$ and $j$, we first searched for evidence that the sites were linked, regardless of the specific alleles involved. To do so, we calculated the total number of shared barcodes $\mathcal{B}_{ij}$ across all 19 timepoints. Under the null hypothesis, $\mathcal{B}_{ij}$ should be Poisson distributed with mean

$$\overline{\mathcal{B}}_{ij} = \sum_t q_{jt} \sum_{\mu \in \mathscr{B}_t} G_t(D_\mu) \,. \tag{20}$$

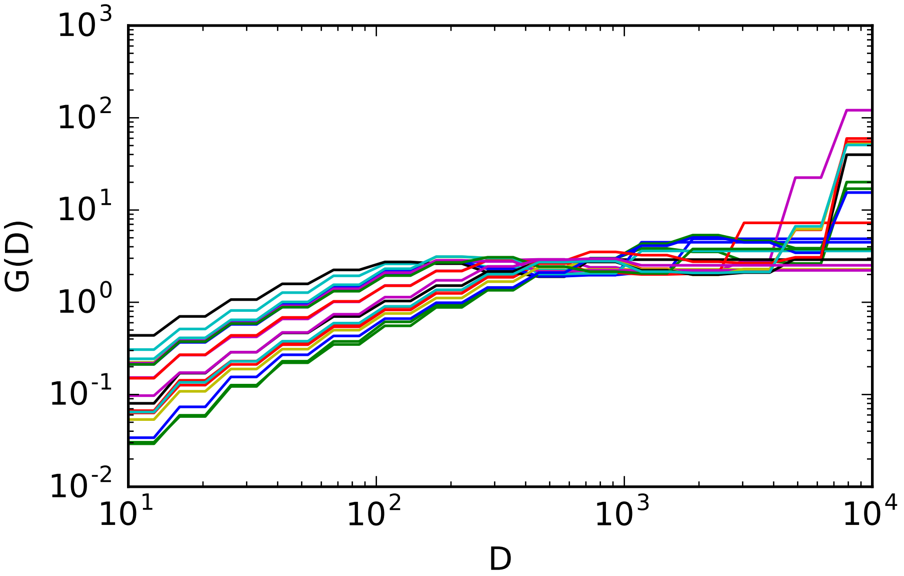

Figure S16: Calibrated null model of read cloud sharing as a function of read cloud coverage. Colored lines denote the $\hat{G}(D)$ function in Eq. (19) estimated for each of the 19 timepoints.

Using the observed value of $\mathcal{B}_{ij}$, we then calculated a $P$-value,

$$P_{ij} = \sum_{k \geq \mathcal{B}_{ij}} \frac{\overline{\mathcal{B}}_{ij}^k e^{-\overline{\mathcal{B}}_{ij}}}{k!}, \tag{21}$$

which quantifies how surprising the observed levels of barcode sharing are under the null hypothesis. We used a Bonferonni correction to control for multiple comparisons, recording a significant hit when the corrected $P$-value was less than 0.1 and the total number of shared read clouds was $\geq 12$.

We applied this approach in two different contexts. In the first case, we scanned for evidence of genetic linkage *within* the subset of temporally varying SNVs identified in each panel of Fig. 3 (Supplemental Data S4). We performed this test separately to maximize statistical power within this subset of highly interesting SNVs. For the second application (Fig. 4), we performed a genome-wide scan for all pairs of SNVs within each of the species in Fig. 2. To preserve statistical power, we again restricted this scan to pairs of SNVs in the same species, since examples of SNV sharing across species are expected to be rare (see Section 7.5).

##### *7.4.2 Quantifying linkage at the allelic level*

For each of the significant SNV pairs identified above, we next sought to quantify the patterns of genetic linkage at the allelic level. For a pair of biallelic sites, there are 4 possible combinations of alleles (or *haplotypes*) that can be observed together in a read cloud. We refer to these using the standard notation, $ab$, $aB$, $Ab$, and $AB$ (Fig. 4A), with $h$ denoting an arbitrary haplotype within this set. As above, these combinations could reflect true examples of linkage on the same DNA fragment, or spurious signals produced by two separate fragments in the same read cloud. To distinguish between these two scenarios, we repeated our null model calculations above, with $i$ and $j$ now ranging over the two alleles at each site. We calculated the total number of shared read clouds ($\mathcal{B}_{ij,h}$) for each $h \in \{ab, aB, Ab, AB\}$, as well as the corresponding expected values ($\overline{\mathcal{B}}_{ij,h}$).

In this case, however, it is important to augment the null hypothesis to include the possibility of sequencing or PCR errors, since an error at the target site can produce a spurious haplotype where none actually exists. We incorporated this additional source of error using the empirical rates estimated in Section 5.4. We first calculated an average error rate,

$$\overline{p}_{\mathrm{err,ij}} = \langle p_{\mathrm{err},i} \rangle_{B.\ vulgatus} \ , \tag{22}$$

where the average is taken over all sites in *Bacteroides vulgatus*. This species was chosen because it has one of the highest coverages and SNV densities across our dataset, yielding the most robust estimates of $p_{\mathrm{err}}, i$. We used this average value to form a pair-specific error estimate,

$$p_{\mathrm{err},ij} = \max\left\{\overline{p}_{\mathrm{err}}, p_{\mathrm{err},i}, p_{\mathrm{err},j}\right\} \ , \tag{23}$$

and we augmented the expected number of shared read clouds by

$$\overline{\mathcal{B}}_{ij,h} \to \overline{\mathcal{B}}_{ij,h} + p_{\mathrm{err},ij}\mathcal{B}_{ij} \ , \tag{24}$$

where $\mathcal{B}_{\mathrm{ij}}$ is the total number of observed read clouds shared between the two SNVs.

Based on this null model, we tested for the presence of the various haplotypes in a specific order, which was designed to maximize statistical power and provide biological insights into the patterns of linkage between the two SNVs. For each species, we first compiled a list of significantly linked SNV pairs using the procedure described above. We only considered SNVs in the personal core genome, and we excluded SNV pairs with $\ell > 50$kb or $p_{\mathrm{err}} > 0.01$. To ensure that there were sufficient opportunities to observe instances of 3 or 4 haplotypes, we only considered SNV pairs where all single-site alleles were present in at least 10% of the shared read clouds, and with at least 4 counts in total. We also restricted our attention to SNV pairs where we expect to observe at least 4 reads for the smallest haplotype under linkage equilibrium ($\mathcal{B}_{ij} f_i f_j \geq 4$). The total number of resulting SNV pairs for each species is shown in Fig. 4C. We then examined the linkage structure within each of these SNV pairs as described below.

1. **Perfect linkage between SNV pairs.** The simplest form of genetic linkage between two SNVs is known as *perfect linkage*. It describes a situation where a mutation at one site always co-occurs with a mutation at the other site (consistent with a mixture of just two strains). In terms of the four haplotypes, perfect linkage requires that either $ab/AB$ are present and $aB/Ab$ are absent, or vice versa. Deviations from this pattern indicate that the two mutations occurred in a nested fashion (or have undergone recombination, see below), and require at least three haplotypes to explain. To scan for deviations from perfect linkage, we tested whether $\mathcal{B}_{ab} + \mathcal{B}_{AB}$ and $\mathcal{B}_{Ab} + \mathcal{B}_{aB}$ were both higher than expected by chance, using the same procedures as described in Eq. (21) above. To correct for multiple hypothesis testing, we converted the raw $P$-values into genome-wide $Q$-values using the formula,

$$Q_{ij} = \min_{Q > P_{ij}} \left\{ \frac{Q \sum_{i',j'} 1}{\sum_{i',j'} \theta(Q - P_{i'j'})} \right\} \ . \tag{25}$$

   rejecting perfect linkage if $Q_{ij} < 0.05$. To be conservative, we only considered SNV pairs with at least 3 read clouds supporting the alternate haplotype pair. We performed this calculation for the entire set of SNV pairs in each species, as well as subsets stratified by different values of $\ell$ (Fig. 4D). Some species are dominated by SNV pairs in perfect linkage, while other species have a significant fraction of SNV pairs supporting 3 or more haplotypes.

2. **The four gamete test and recombination.** SNV pairs that are not in perfect linkage can arise in two ways. They can be produced by nested mutation events, where only a subset of the individuals with the first mutation will also possess the second one. This scenario is often referred to as *complete linkage*, since it consistent with a single origin of each mutation and purely clonal inheritance. Alternatively, violations of perfect linkage can arise from recurrent mutations or recombination between the two SNVs. In the limit of infinite recombination, the haplotypes frequencies should approach *linkage equilibrium*,

$$\mathcal{B}_{ij} \approx \frac{\left(\sum_j \mathcal{B}_{ij}\right)\left(\sum_i \mathcal{B}_{ij}\right)}{\sum_{ij} \mathcal{B}_{ij}}\,, \tag{26}$$

which is equivalent to randomly permuting the observed alleles across the sampled read clouds. Complete linkage can be ruled out if all four haplotypes are present in the sample (the so-called *four gamete test* [18]). We implemented a version of the four gamete test using our error model above, by testing whether the $\mathcal{B}_{ij,h}$ values for all four haplotypes were higher than expected by chance. Four haplotypes were deemed to be present if the adjusted $Q$-value was less than 0.05, and if there were at least 3 read clouds supporting the smallest haplotype; all other SNVs pairs were classified as being in complete linkage. We performed this calculation for the entire set of SNV pairs where perfect linkage could be rejected, as well as subsets stratified by different values of $\ell$ (Fig. 4D). Only a small fraction of 4-haplotype SNVs are observed across the different species. Furthermore, this fraction remains constant (or even decreases) for SNVs separated by increasing genomic distances, in contrast to the recombination-mediated decay of linkage observed in the dominant strains from different hosts [4]. This provides further evidence that within-host recombination events are rare in our dataset.

#### 7.5 Inferring linkage between SNVs and species backbones

In addition to inferring linkage between specific pairs of SNVs, we also used the read cloud information to determine whether the temporally variable SNVs in Fig. 2C were linked to the correct species backbone. To do so, we leveraged patterns of read cloud sharing between the temporally variable SNVs and the core genomes of each of the species in our personalized reference panel. Because these core genes were defined to be present in most strains of a given species, we hypothesized that they would be a useful proxy for the true species backbone. For each of the SNVs in Fig. 2C, we created a list of the core genes that shared read clouds with either of the two alleles, and we also kept track of the number of shared read clouds for each allele/gene pair. For species with more than 1000 temporally variable SNVs, we chose a random subset of 1000 SNVs to serve as a representative sample. For each allele, we recorded the 10 most frequently shared genes, provided that they shared at least 5 read clouds in total and were present in least 5% of the read clouds that mapped to that allele. We then calculated the fraction of these top scoring genes that originated from the correct species. We considered there to be a positive confirmation if, for both alleles, more than two thirds of the top genes originated from the correct species. We considered there to be a negative confirmation if either of the alleles had more than a third of their top genes originating from a different species.

The results are shown in Supplemental Fig. S9. Across species, we observe an average positive confirmation rate of $\approx 80\%$ and a negative confirmation rate of $< 1\%$, with most of the negatives clustering in just 3 species (where they still comprise only a small fraction of the total identified SNVs). This is an important sanity check: it suggests that many of the temporally variable SNVs identified in Section 5.5.1 are truly changing within their respective species, and are not solely driven by mis-mapped reads from other temporally fluctuating species.

### 8 Clustering SNVs into haplotypes

The low levels of recombination in Fig. 4 suggest an interesting simplification, in which the dynamics of many individual SNVs may be captured by a mixture of just a few clonal haplotypes (or "strains"). Several approaches have been developed to infer the sequences of these haplotypes and their mixture proportions using the co-occurrence patterns of SNVs in different timepoints [19–22]. However, since SNV frequencies can always be fit exactly using a sufficiently large number of haplotypes, these methods suffer from an inherent overfitting problem, and it is necessary to constrain both the number and structure of the haplotypes to be fitted. This is typically done by first fixing the number of haplotypes at some value $K$, and then using model selection criteria like AIC [20, 22], Bayes factors [19], or other heuristic approaches [21] to choose the optimal value of $K$. While these approaches have been shown to work well in certain test cases, they can be difficult to extrapolate to new datasets, since it is not always clear how the tradeoff between model fit and model simplicity interacts with the geometry of haplotype space. For similar reasons, it is not known how the presence of a small fraction of outlier SNVs (e.g. from mapping errors or other metagenomic artifacts) could influence the reported results.

To illustrate these issues more concretely, we focused on the *Bacteroides vulgatus* population in Supplemental Fig. S10. This species has one of the highest rates of intra-population diversity across our panel (Fig. 2B) while also displaying high levels of linkage disequilibrium (Fig. 4D), making it an ideal candidate for compression via haplotype clustering. In addition, this species benefits from one of the highest levels of sequencing coverage across our panel, making it a best-case scenario for existing strain detection algorithms that leverage temporal correlations in SNV frequencies.

We used the recently published StrainFinder [22] program as a representative example of existing algorithms. We ran StrainFinder (using default parameters) on the collection of ∼16,000 SNVs in the resident genome that reached intermediate frequency ($0.2 < f < 0.8$) in at least one timepoint. The optimal AIC score was observed for $K = 9$ strains, after scanning over values in the range $2 \leq K \leq 20$. The estimated frequencies of these 9 strains, along with the predicted and observed SNV frequencies, are shown in Supplemental Fig. S17. The estimated strain frequencies seem reasonable at first glance, but closer examination of the predicted SNV trajectories reveals a peculiar banded structure, suggesting that StrainFinder is going to great lengths to fit some other systematic deviation from the model, and that the inferred haplotypes may not be trustworthy. To test this hypothesis, we used the 9 estimated strain genotypes to generate predictions for linkage disequilibrium between pairs of SNVs, similar to what we measured in Fig. 4C,D. We observed violations of the 4-gamete test among these 9 predicted strains for ∼10% of the SNV pairs with $\ell$<50kb. This is more than an order of magnitude higher than the rate estimated from the read cloud data in Fig. 4D, and suggests that the haplotypes estimated by StrainFinder may contain significant errors. Since we expect *B. vulgatus* to be a best-case scenario for this program (and that there will be reduced power to detect similar errors in other species), we decided not to apply these existing strain detection methods in the present study.

Instead, we pursued a non-parametric approach inspired by recent experimental evolution studies [9, 11], which leverages the graphical information contained in joint allele frequency trajectories like Fig. 3. Rather than aiming to reconstruct the full set of haplotypes, the goal of this approach is to first identify clusters of SNVs that are in perfect LD with each other, so that they can be replaced with a single representative trajectory. In simple cases, coarse-grained features of the

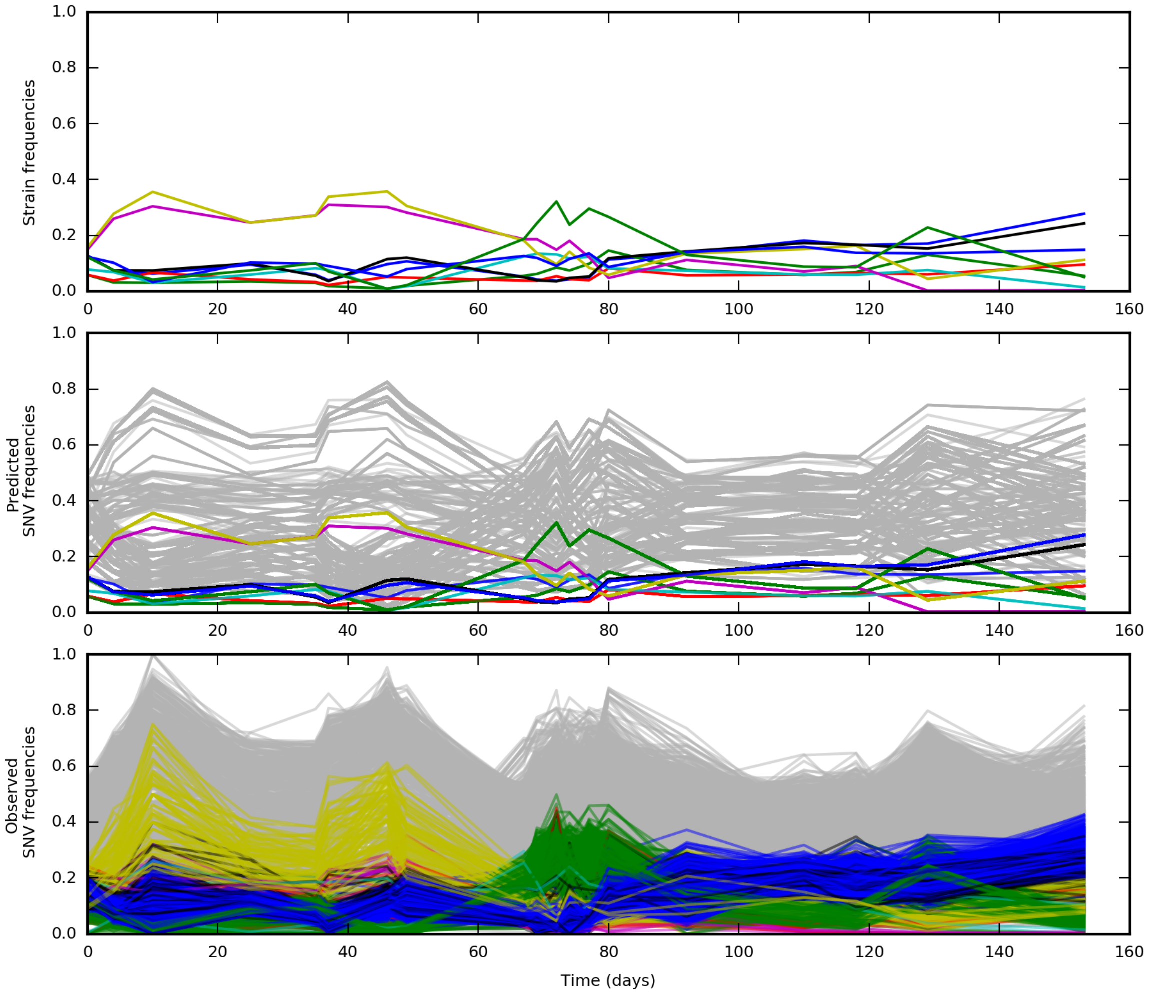

Figure S17: Results from the StrainFinder algorithm [22] for the *Bacteroides vulgatus* population. The top panel shows the estimated frequencies for each of the 9 predicted strains. The middle panel shows the expected SNV frequencies obtained by combining the strain frequencies with the inferred strain genotypes. The bottom panel shows the observed SNV frequencies.

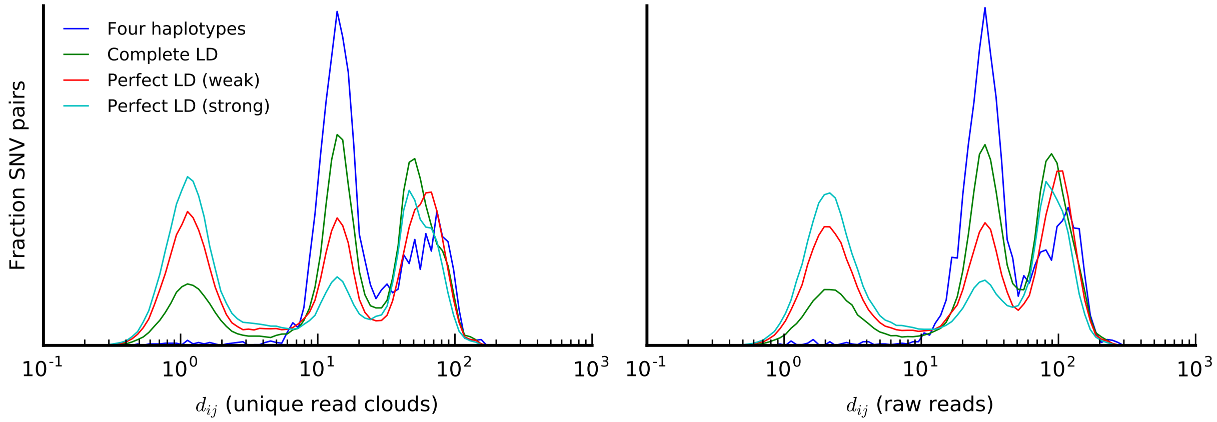

Figure S18: The distribution of the distance metric $d_{ij}$ for all pairs of *Bacteroides vulgatus* SNVs in Supplemental Fig. S17. SNV pairs are classified according to the linkage categories using read cloud information as described in Section 7.4. We have further divided the perfect LD category into weak and strong sub-classes, respectively, depending on whether we observe any read clouds that support the third haplotype.

haplotype dynamics can be visually inferred from the dynamics of the relatively small number of cluster trajectories (e.g. Fig. 3 in Ref [9] or Fig. 5 in Ref [23]), while systematic errors and other metagenomic artifacts can be identified and excluded separately (see, e.g. Ref [11]).

We developed an agglomerative clustering method to identify SNV clusters based on similarities in their allele frequency trajectories. We used a distance metric informed by binomial sampling noise expected for allele frequency data. For two observed trajectories $\hat{f}_{it}$ and $\hat{f}_{jt}$ with the same relative polarization, we defined the distance metric,

$$d(\hat{f}_{it}, \hat{f}_{jt}) = \frac{1}{T}\sum_{t=1}^{T} \frac{2(\mathcal{D}_{it}+\mathcal{D}_{jt})(\hat{f}_{it}-\hat{f}_{jt})^2}{\left(\hat{f}_{it}+\hat{f}_{jt}\right)\left(1-\hat{f}_{it}+1-\hat{f}_{jt}\right)}\,, \tag{27}$$

where $D_{it}$ and $D_{ij}$ are the total sequencing coverages at each timepoint. If the true frequencies were identical, we would expect the product $T \cdot d(\hat{f}_{it}, \hat{f}_{jt})$ to approach a $\chi^2_T$ distribution in the limit of large $D$, so that the typical distances should be of order $d \approx 1 \pm 2/\sqrt{T}$.

However, our task is complicated by the fact that we do not typically know the relative orientations of SNVs in natural microbiome populations. To overcome this problem, we defined a pair of distance matrices,

$$\begin{aligned} d^{+}_{ij} &= d(\hat{f}_{it}, \hat{f}_{jt})\,, \\ d^{-}_{ij} &= d(\hat{f}_{it}, 1-\hat{f}_{jt})\,, \end{aligned} \tag{28}$$

that account for both relative polarizations of each SNV pair, along with an effective distance,

$$d_{ij} = \min\left\{d^{+}_{ij}, d^{-}_{ij}\right\}\,. \tag{29}$$

Under perfect linkage, we expect that the smaller value of $d^{+}_{ij}$ and $d^{-}_{ij}$ will typically indicate the correct relative polarization, so that the typical value of $d_{ij}$ should still be close to one.

We performed UPGMA clustering on the distance matrix $d_{ij}$ using the `linkage` function in the SciPy hierarchical clustering package [24], and we used a maximum distance criterion to identify flat SNV clusters from the resulting UPGMA dendrogram. We used an empirically calibrated distance threshold derived from the patterns of read cloud sharing in *B. vulgatus*. Supplemental Fig. S18 shows the distribution of distances $d_{ij}$ for pairs of SNVs in Supplemental Fig. S17 with significant rates of read cloud sharing, stratified according to the linkage categories described in Section 7.4. As expected, SNVs that are observed to be in perfect linkage have a mode near $d_{ij} \approx 1$, consistent with our theoretical predictions above. However, we also observe two additional modes at much larger distances, similar to SNV pairs in complete LD or where all four haplotypes are present. We assumed that the first mode reflects SNV pairs that are truly in perfect LD, while the second two modes reflect mis-classified SNV pairs produced by finite sampling effects (e.g., not observing the third haplotype in a small sample of read clouds) or within-population variation in the genomic locations of different SNVs. In either case, this finding suggests that naive clustering based on read cloud sharing alone is likely to yield a substantial fraction of false positives. Based on this data, we estimated a distance threshold of $d^* \approx 3.5$ that separates the perfect linkage mode from the modes at higher distances. We used this threshold to form flat SNV clusters using `fcluster` function in SciPy [24], and we used the relative values of $d^-_{ij}$ and $d^+_{ij}$ to determine the relative polarization of the SNVs in each cluster. We then estimated a frequency trajectory for each cluster by summing up the ancestral and derived read counts of the polarized SNVs.

When applied to *B. vulgatus*, this methods suggests that ≈85% of the SNVs can be grouped into just 3 SNV clusters, which persist at intermediate frequencies throughout the sampling period (Supplemental Fig. S10). The trajectories of the three clusters suggest a mixture of three dominant haplotypes: (i) the ancestral alleles of all three clusters, (ii) the derived alleles of the blue and green clusters and the ancestral allele of the orange cluster, and (iii) the derived alleles of the green and orange clusters and the ancestral allele of the blue cluster. In this case, the three SNV clusters contain a sufficiently large number of SNVs that we can test this hypothesis directly using the patterns of read cloud sharing between SNVs in each cluster (Supplemental Fig. S10, Supplemental Data S7). This analysis confirms that each of the clusters is consistent with perfect LD, while the linkage across clusters was consistent with the haplotype structure above. This provides a striking example of the oligo-colonization model suggested by Ref [4] and others, in which just a few distantly related strains can be maintained at intermediate frequencies in the same host.

We used this same approach to cluster the temporally variable SNVs in Fig. 3. To boost computational efficiency for species with large numbers of SNVs, we used a modified version of UPGMA clustering in which flat clusters are seeded using a random subset of 1000 of the temporally variable SNVs, and the remaining SNVs are assigned to those clusters if the distance between the cluster trajectory and the SNV trajectory was $< d^*$. This results in a 10-fold speedup for species like *E. eligens* and *B. vulgatus*, with minimal loss of accuracy.

### 9 Distinguishing between genetic drift and natural selection in allele frequency time series

To show that the allele frequency changes in Fig. 3 are inconsistent with a simple model of genetic drift, we developed a new statistical test to compare these data against the standard Wright-Fisher diffusion,

$$\frac{\partial f}{\partial t} = \sqrt{\frac{f(1-f)}{N_e}}\eta(t)\,, \tag{30}$$

where $N_e$ is an effective parameter describing the (constant) strength of genetic drift and $\eta(t)$ is a Brownian noise term. Note that $f(t)$ here is the true allele frequency, rather than the sample frequency $\hat{f}_t$ estimated from the read counts. The two are connected through an additional sampling process,

$$\mathcal{A}_t \sim \text{Binomial}(\mathcal{D}_t, f(t)) \,, \tag{31}$$

where $\mathcal{A}_t$ and $\mathcal{D}_t$ respectively denote the alternate allele count and total coverage at timepoint $t$. at the observed coverage $\mathcal{D}_{it}$. Together, Eqs. (30) and (31) constitute a hidden Markov model for the observed frequency trajectory with two unknown parameters: the effective population size $N_e$ and the value of $f(t)$ at some reference timepoint $t_0$ (discussed in more detail below). The goal is to determine when a single realization of this stochastic process is sufficient to rule out the general class of models described by Eq. (30).

Several previous approaches have been developed to address this problem [25–30]. The vast majority of these methods attempt to reject the model in Eq. (30) by comparing it to a model with an explicit natural selection term,

$$\frac{\partial f}{\partial t} = s_e f(1-f) + \sqrt{\frac{f(1-f)}{N_e}}\eta(t) \,, \tag{32}$$

with a constant effective selection coefficient $s_e$. In this case, rejecting the neutral model in Eq. (30) is equivalent to showing that $s_e$ is significantly different from zero.

However, in large microbial populations, the natural selection term may behave very differently from the simple form assumed in Eq. (32). For example, genetic linkage between selected mutations will generally cause the selection coefficient to take on a time dependent form,

$$\frac{\partial f}{\partial t} = s_e(t) f(1-f) + \sqrt{\frac{f(1-f)}{N_e}}\eta(t) \,, \tag{33}$$

where $s_e(t)$ is the difference between the mean fitness of all the haplotypes that cary the alternate allele and the mean fitness of the haplotypes that carry the reference allele. The effective selection coefficient in Eq. (33) may only be tangentially related to the direct selection pressure on the allele itself, since it emerges from a sum of selection pressures across many genomic loci. Since the frequencies of the underlying haplotypes will generally shift over time, the effective selection coefficient can be time-dependent even when the fitnesses of the underlying haplotypes are fixed. Furthermore, we note that our present study focuses on a highly non-equilibrium scenario involving the transient introduction and removal of antibiotics, as well as the correlated perturbations this induces in the species level composition of the community. It is therefore reasonable to suppose that selection pressures will vary over time due to these external reasons as well. We therefore require methods for distinguishing Eq. (30) from the more general class of selection models encoded in Eq. (33).

A simple extension of the existing parametric approaches proves to be difficult, since Eq. (33) is capable of fitting any frequency trajectory exactly with the appropriate choice of $s_e(t)$. We therefore developed a goodness-of-fit test to directly quantify how unlikely the observed data are under the model in Eq. (30), without explicitly comparing to Eq. (33).

The basic idea behind our approach is simple: though there is considerable freedom to generate different trajectories under the random process in Eq. (30), these trajectories must still satisfy several statistical regularities. For example, genetic drift requires a time of order $\Delta t \sim N_e$ to shift the allele frequency from low frequencies (e.g. 20%) to high frequencies (e.g. 80%). If we observe a shift in frequency of this magnitude over a time interval $\Delta t$, then we know that $N_e \sim \Delta t$, and the corresponding shifts in frequency across all the other time intervals in the trajectory must be consistent with this estimate. In particular, there is a high probability that the mutation will have either fixed or gone extinct after another $\Delta t$ generations. Thus, a characteristic signal against the constant genetic drift model in Eq. (30) arises when the frequency changes by a large amount in some time intervals, and then changes more slowly in others, particularly if the allele is close to one of the boundaries at the later timepoints.

Our present method is an attempt to make this intuition precise, while accounting for the additional complications of sampling noise and the fact that we have conditioned on observing at least one large frequency change in scanning for temporally variable SNVs in Section 5.5.1. As a goodness-of-fit statistic, we used likelihood of the data under the best fit model from the class in Eq. (30):

$$\Lambda(\{\mathcal{A}_t\}) = -\log \Pr[\{\mathcal{A}_t\}|\{\mathcal{D}_t\}, \hat{N}_e] \,, \tag{34}$$

where $\hat{N}_e$ is the best-fit value of $N_e$.

Since we are working with a hidden Markov model, the log-likelihood can be calculated efficiently using standard dynamic programming approaches. In particular, the backward table is defined by the recursion relation,

$$B(f, t_{k-1}) = \int df' p(f'|f, \Delta t = t_k - t_{k-1}, N_e) \cdot \Pr(\mathcal{A}_{t_k}|\mathcal{D}_{t_k}, f') \cdot B(f', t_k) \,, \tag{35}$$

where $\Pr(\mathcal{A}|\mathcal{D}, f)$ is the binomial sampling distribution in Eq. (31) and $p(f'|f, \Delta t, N_e)$ is the transition probability defined by the diffusion process in Eq. (30). We calculated this transition probability by exploiting the known equivalent between the diffusion process and discrete individual Wright-Fisher model at a finite value of $N_e$:

$$p(f'|f, \Delta t = 1, N_e) = \binom{N_e}{N_e f'} f^{N_e f'} (1-f)^{N_e(1-f')} \,, \quad f, f' = 0, \frac{1}{N_e}, \ldots, \frac{N_e - 1}{N_e}, 1. \tag{36}$$

Extensions to $\Delta t > 1$ can be obtained by simple matrix multiplication. The likelihood follows from integrating the backward table over the initial frequency

$$\Pr[\{\mathcal{A}_t\}|\{\mathcal{D}_t\}, N_e] = \int df p_0(f_0) B(f_0, t_0) \,, \tag{37}$$

where $p_0(f_0)$ is a prior distribution described in more detail below.

To account for the additional conditioning caused our scan for temporally variable SNVs, we split the trajectory into two smaller segments. The latter segment contains all the timepoints after the sample frequency first exceeded 70%, while the prior segment contains all the timepoints before this point from the initial timepoint until the sample frequency was last below 20%. Any timepoints that fall between the two segments (typically the antibiotic timepoints) were excluded from further analysis. We then approximated the likelihood of the full trajectory as the product between independent trajectories for the later and prior segments:

$$\Pr[\{\mathcal{A}_t\}|\{\mathcal{D}\}, N_e] = \Pr[\{\mathcal{A}_t\}_{\text{prior}}|\{\mathcal{D}\}, N_e] \cdot \Pr[\{\mathcal{A}_t\}_{\text{latter}}|\{\mathcal{D}\}, N_e] \,. \tag{38}$$

The likelihood of the latter segment can be calculated from the algorithm described above, while the likelihood of the prior segment can be calculated from the same algorithm using the time-reversed version of the trajectory, where the final timepoint of the prior segment is used as the initial timepoint of the time-reversed process.

Using this approach, we evaluated the likelihood of a given trajectory across a grid of 11 logarithmically spaced $N_e$ values, and we identified the maximum likelihood estimator,

$$\hat{N}_e = \mathrm{argmax}_{N_e} \Pr[\{\mathcal{A}_t\}|\{\mathcal{D}\}, N_e] \, . \tag{39}$$

Since the "initial" timepoints for the likelihood calculation were chosen to be in the interior of the trajectory, we used a uniform prior for $f_0$ in Eq. (37). Using the inferred value of $\hat{N}_e$, we simulated $n = 10,000$ bootstrapped trajectories from the null model in Eq. (30). For each trajectory, we calculated the goodness-of-fit statistic $\Lambda$ in Eq. (34), and we compared it to the observed value of $\Lambda$ calculated from the data. We then defined a $P$-value based on the fraction of bootstrapped samples that were more extreme than the corresponding value of $\Lambda$ for the observed trajectory:

$$P = \Pr[\Lambda \geq \Lambda_{\mathrm{obs}}] \approx \frac{1 + \sum_{\ell=1}^{n} \theta(\Lambda_\ell - \Lambda_{\mathrm{obs}})}{n+1} \, . \tag{40}$$

We applied this approach to the temporally variable SNV trajectories from the example species in Fig. 3. To minimize the effects of sampling noise, we we merged the alternate and reference read counts from each SNV cluster into a single effective trajectory (up to a maximum coverage of $D = 2000$ in any individual timepoint). We carried out this procedure for the full timecourse, as well as a second version where we fit a different value of $N_e$ separately for each of the two segments. The $P$-values for the combined timecourse, as well as for the individual segments are shown in Supplemental Table S3. The neutral model in Eq. (30) provides a poor fit to the observed data for all species except *A. finegoldii*.

Supplemental Table S1: List of samples and associated metadata.

Supplemental Table S2: DNA yield and quality from different extraction methods.

Supplemental Table S3: Evidence for natural selection in SNV frequency trajectories. Results of the natural selection test in Section 9 for each of the species in Fig. 3.

### Supplemental Data Files

***Supplemental Data S1***

List of SNV differences shown in colored lines in Fig. 3. SNV clusters with more than 300 SNVs (e.g. *E. eligens*) were randomly downsampled to ∼300 SNVs.

***Supplemental Data S2***

Analogous version of Supplemental Data S1 for Supplemental Fig. S7.

***Supplemental Data S3***

Analogous version of Supplemental Data S1 for Supplemental Fig. S8.

***Supplemental Data S4***

Statistics of read cloud sharing between SNV differences in Fig. 3.

***Supplemental Data S5***

Fragment analyzer output for HMW DNA samples used in this study (Part 1/2). Each page shows the size distribution of HMW DNA extracted from different study timepoints (page 4-15), as well as the standard ladder (page 16), using the sample ids in Supplemental Table S1.

***Supplemental Data S6***

Fragment analyzer output for HMW DNA samples used in this study (Part 2/2).

***Supplemental Data S7***

Aggregate patterns of read cloud sharing for each of the SNV clusters in the *B. vulgatus* example in Supplemental Fig. S10.

[23] Shijie Zhao, Tami D Lieberman, Mathilde Poyet, Mathieu Groussin, Sean M Gibbons, Ramnik J Xavier, and Eric J Alm. Adaptive evolution within the gut microbiome of individual people. *bioRxiv*, page 208009, 2017.
